# Interplay of YEATS2 and GCDH regulates histone crotonylation and drives EMT in head and neck cancer

**DOI:** 10.1101/2024.09.24.614679

**Authors:** Deepak Pant, Parik Kakani, Rushikesh Joshi, Abin Sabu, Shruti Agrawal, Atul Samaiya, Sanjeev Shukla

**Affiliations:** Department of Biological Sciences, Indian Institute of Science Education and Research Bhopal, Bhopal, Madhya Pradesh 462066, India; Department of Pathology, Bansal Hospital, Bhopal, Madhya Pradesh 462016, India; Department of Surgical Oncology, Bansal Hospital, Bhopal, Madhya Pradesh 462016, India

**Keywords:** Head and neck cancer, epigenetics, YEATS2, histone crotonylation, GCDH, EMT

## Abstract

The regulation of gene expression is an integral cellular process orchestrated by epigenetic marks like histone modifications. Perturbations in the activity or abundance of epigenetic factors can lead to tumorigenesis. Remarkably, several metabolites influence the epigenetic landscape in cells. Here, we investigated the interplay between a highly expressed epigenetic factor, YEATS2, and a metabolic enzyme, GCDH, in regulating epithelial-to-mesenchymal transition in head and neck cancer. We report that the histone reader YEATS2 is responsible for increased invasive potential in head and neck cancer in an SP1-dependent manner. YEATS2 functions by maintaining histone crotonylation, and its abrogation leads to a global decrease in the H3K27cr mark. Mechanistically, we report that YEATS2 maintains high promoter H3K27cr levels by assisting in the recruitment of crotonyltransferase p300 at the promoter of the EMT-promoting gene *SPARC*. Further, we found that the addition of the H3K27cr mark is also dependent on the crotonyl-CoA-producing enzyme GCDH. Overall, we describe a novel mechanism of interplay between epigenetics and metabolism in head and neck tumorigenesis, which results in the enhanced expression of EMT-related genes in a histone crotonylation-dependent manner.

## Introduction

Head and neck cancer is an umbrella term associated with a number of cancers of the oral cavity and pharynx. Like other solid tumors of epithelial origin, head and neck cancer (HNC) involves a multi-step progression towards malignancy, of which transition from an epithelial to a mesenchymal phenotype is a pivotal event ^1^. This epithelial-to-mesenchymal transition (EMT) involves dramatic changes like loss of cell polarity and acquisition of migratory and invasive properties in the tumor cells ^2^. Underlying these remarkable external changes in tumor cells, are equally notable changes in gene expression and signalling pathways. Moreover, widespread alteration in the gene expression program of cancer cells undergoing EMT can be a result of either altered activity or the change in abundance of several epigenetic factors. Tumor-specific epigenomic changes thus allow HNC cells to tweak the expression levels of multiple genes at once, thereby leading to a drastic transformation in phenotype during metastasis. These large-scale changes are due to the rapid and global nature of epigenetic alterations ^3^.

Similar to epigenetic changes, metabolic rewiring is another important hallmark of cancer cells capable of carrying out vast tumor-promoting changes rapidly. Numerous reports in the past have documented instances of loss-of-function mutations or changes in the gene expression of enzymes involved in key metabolic processes, such as glycolysis or the citric acid cycle, to promote oncogenesis ^4^. However, the discovery of non-canonical functions of several enzymes and their atypical localization in the nucleus has intrigued cancer researchers of late. Metabolic enzymes are now believed to play a more direct role in influencing oncogene expression via crosstalk with the epigenome. One such phenomenon is the role of metabolites produced as a result of aberrant metabolism in acting as substrates for epigenetic factors. A ground breaking study in this regard by Wellen et al. reported the nuclear localization of a mitochondria-resident protein ACLY/ACL (adenosine triphosphate–citrate lyase). Nuclear pool of ACLY was found to affect histone acetylation directly by producing acetyl-coenzyme A (CoA) in colon carcinoma cells ^5^. Moreover, another study reported a direct interaction involving α-ketoglutarate dehydrogenase (α-KGDH) and a histone acetyltransferase leading to the production of localized succinyl-CoA in nucleus, which in turn led to histone succinylation and enhanced expression of genes related to tumor cell proliferation ^6^. M2 isoform of pyruvate kinase (PKM2) is another metabolic enzyme known to exhibit a non-canonical function by directly affecting gene transcription. We have recently reported the role of nuclear PKM2-HIF1α-p300 axis in maintaining high levels of H3K9ac marks at the promoter of *PFKFB3*, which thereby leads to its increased expression in hypoxic breast cancer cells ^7^. The reports stated herein, along with others ^8^, delineate the role of the nexus between aberrant metabolism and the epigenome in regulating various aspects of tumorigenesis in a wide variety of cancers. However, the significance of this partnership in regulating EMT, which is one of the most vital events in cancer, remains largely unexplored.

In this study, we have made an attempt to dissect the role of previously unknown link between epigenetic factors and metabolism in promoting EMT in head and neck cancer. To explore tumor-promoting epigenetic factors in HNC we analyzed large gene expression dataset and narrowed down our search to YEATS2. This gene was found to be overexpressed in HNC, had significant correlation with EMT, and also showed association with poor patient prognosis. YEATS2 (YEATS-domain containing protein 2) is a histone reader protein capable of recognizing epigenetic marks such as histone acetylation and crotonylation ^9^. Through RNA-seq, it was revealed that silencing of YEATS2 led to a decrease in the expression of multiple EMT-associated genes. Further, the transcription factor SP1 was found to be responsible for YEATS2 expression in HNC. While exploring the relationship of YEATS2 with cancer cell metabolism, we found that YEATS2 is co-expressed with crotonyl-CoA producing GCDH (glutaryl-CoA dehydrogenase) in HNC, and the downregulation of either of these two genes led to a decrease in H3K27cr levels in HNC cells. The H3K27cr mark has been previously reported to activate the transcription of pro-tumorigenic *ETS1* in colorectal cancer ^10^. Corroborating with the previous finding, high H3K27cr levels at the promoter region was associated with an increase in the expression of EMT-related *SPARC* gene. We also demonstrated that YEATS2 interacts with the epigenetic writer p300 to recruit it to the *SPARC* promoter, facilitating the deposition of H3K27cr marks. Subsequent investigation revealed that YEATS2 and GCDH worked synergistically in maintaining higher levels of H3K27cr at the promoter of *SPARC*, thereby causing an increase in its expression. Finally, H3K27cr ChIP-seq in YEATS2-abrogated cells uncovered the role of YEATS2 in maintaining global histone crotonylation on the promoter of a large number of genes, many of those genes being associated with EMT. Altogether, we have elucidated a novel link that unifies epigenetic regulation and metabolic rewiring in contributing toward the enhancement of EMT in head and neck cancer.

## Results

### YEATS2 is overexpressed in head and neck cancer

In order to study cancer-associated epigenetic factors, we analysed publicly available The Cancer Genome Atlas (TCGA) mRNA expression dataset. The analysis aimed at obtaining genes coding for epigenetic factors that are overexpressed in HNC, and are also associated significantly to overall patient survival (**Figure 1A**). The analysis resulted in a list of 10 epigenetic factors that are upregulated in HNC and whose expression levels are associated with poor overall patient survival (**Figure 1B**, **Supplementary Table 1**). Out of these 10 epigenetic factors, *YEATS2*, *RUVBL1* and, *MRGBP* were not explored rigorously in HNC prior to our study ^11–17^. The expression of these three epigenetic factors was thus evaluated at mRNA level using reverse-transcription quantitative real-time PCR (RT-qPCR) in pairs of 8 head and neck normal and cancer tissues collected as a part of this study. We found that *YEATS2,* that encodes for a histone reader protein, was the only gene that was significantly upregulated in cancer as compared to normal tissues (**Figure 1C-D**, **Figure 1—figure supplement 1A-B**). *YEATS2* was also found to be upregulated in publicly available HNC microarray datasets GSE30784 and GSE9844 (**Figure 1E-F**). The association of YEATS2 with EMT was then established by significant positive correlation observed between its expression and the expression of genes included in an EMT gene signature in TCGA data (**Figure 1—figure supplement 1C**). Moreover, the expression of *YEATS2* was upregulated in higher tumor grades (grades 2-4) as compared to grade 1 tumors (**Figure 1— figure supplement 1D**). Additionally, on performing gene set enrichment analysis (GSEA) using TCGA data, we found that YEATS2 expression is significantly correlated with the expression of metastasis-associated genes (**Figure 1—figure supplement 1E**). On further investigation using immunoblotting, we found that the expression of YEATS2 protein was increased in a set of 23 head and neck cancer vs. matched normal tissues (**Figure 1G**, **Figure 1—figure supplement 1F**).

**Figure 1.**
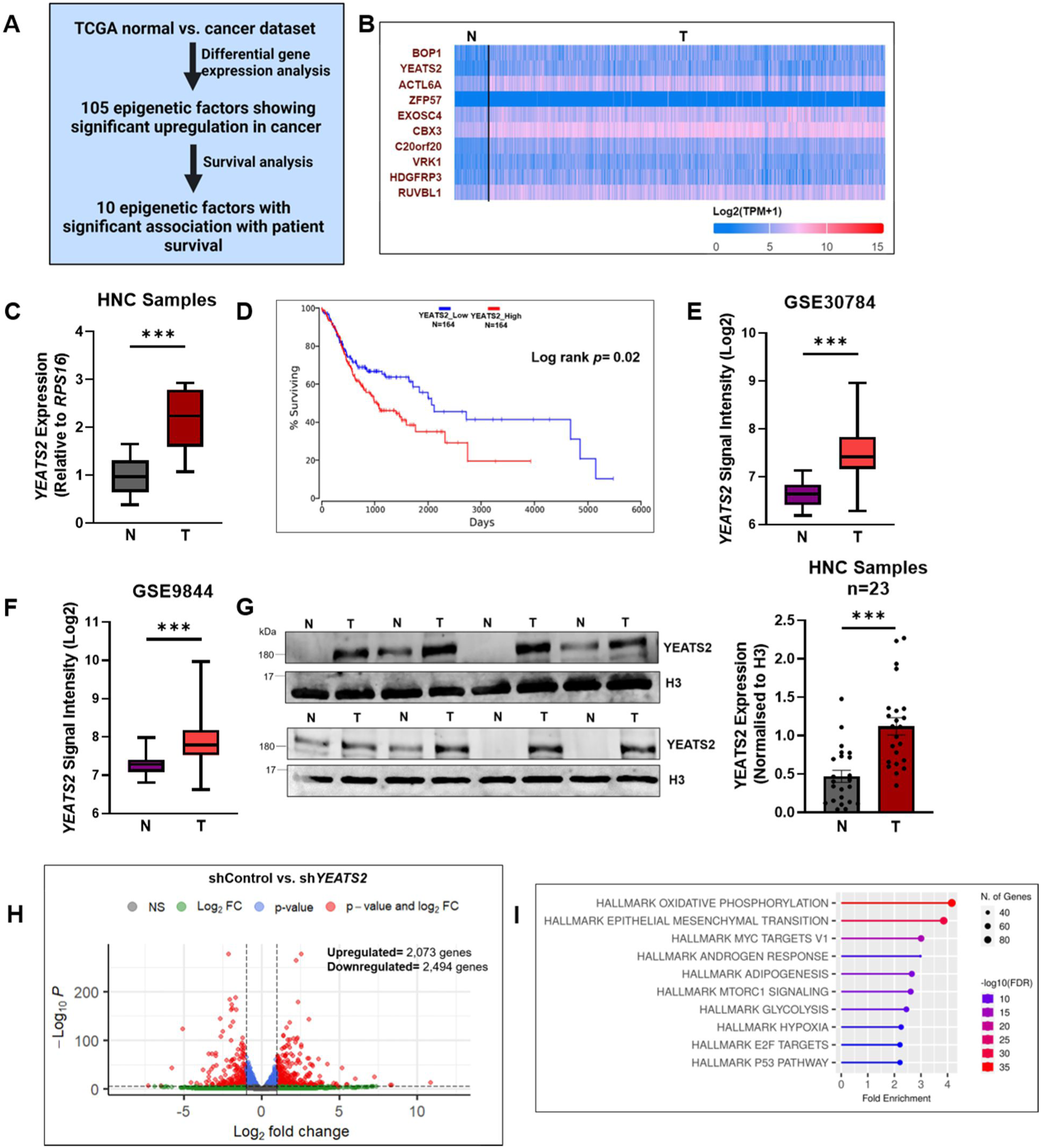
YEATS2 is upregulated in head and neck cancer. **(A)** Computational strategy used to find epigenetic factors with poor patient prognosis overexpressed in HNC. **(B)** Heatmap depicting the TCGA mRNA expression of top 10 epigenetic factors shortlisted using strategy mentioned in (A). **(C)** RT-qPCR result showing mRNA expression of *YEATS2* in HNC samples (n=8). **(D)** Kaplan-Meier curve of *YEATS2* showing association with overall head and neck cancer patient survival. **(E-F)** Gene expression profile of *YEATS2* in publicly available HNC microarray datasets, GSE30784 (E) and GSE9844 (F). **(G)** Immunoblot showing protein levels of YEATS2 in nuclear protein lysates extracted from HNC tissues (n=23). **(H)** Volcano plot of differentially expressed genes in RNA-seq analysis of shControl vs. sh*YEATS2* in BICR10 cells. **(I)** Results of overrepresentation analysis of genes significantly downregulated in shControl vs. sh*YEATS2* RNA-seq data. Error bars, min to max; two-tailed t test, ∗∗∗*p* < 0.001, N-Normal, T-Tumor.

In order to study the role of YEATS2 in the progression of HNC, we performed RNA-seq in YEATS2-silenced BICR10 cells. The shRNA-mediated silencing of YEATS2 led to the downregulation of 2,494 genes, whereas 2,073 genes were found to be upregulated (**Figure 1H, Supplementary Table 2**). Overrepresentation analysis revealed that hallmark gene sets from MSigDb (Molecular signature database) like OXIDATIVE PHOSPHORYLATION and EPITHELIAL MESENCHYMAL TRANSITION were some of the top pathways enriched among the downregulated genes (**Figure 1I**). On the other hand, pathways like TNFA SIGNALING VIA NFKB and G2M CHECKPOINT were enriched among the upregulated genes (**Figure 1— figure supplement 1G**). These set of results hint towards the role of YEATS2 in being one of the essential epigenetic factors responsible for HNC tumorigenesis, mediated via global control of EMT genes’ expression.

### YEATS2 is associated with increased invasion in head and neck cancer cells

Since YEATS2 was found to influence the expression of EMT-related genes globally in our RNA-seq data, we examined the expression of EMT markers in shControl vs. sh*YEATS2* BICR10 and SCC9 cells using immunoblotting and found that Twist1, N-cadherin and, Vimentin showed a decrease upon YEATS2 knockdown (**Figure 2A**, **Figure 2—figure supplement 1A)**. Also, the number of invading cells significantly reduced in Matrigel invasion assay on YEATS2 downregulation, indicating that YEATS2 could be playing a role in the maintenance of EMT phenotype in HNC cells (**Figure 2B**, **Figure 2—figure supplement 1B**). Conversely, the expression of Twist1 was increased and there was significant upregulation in invasive capacity of cells when BICR10 and SCC9 cells were transfected with YEATS2-overexpression construct (**Figure 2C**, **Figure 2—figure supplement 1C-D**). Wound healing assay was performed to assess the cell migration property of HNC cells and it was found that YEATS2-silenced cells migrated slower than control cells (**Figure 2D**, **Figure 2—figure supplement 1E**). On the other hand, overexpressing YEATS2 in BICR10 and SCC9 cells resulted in faster wound healing as compared to control cells (**Figure 2E**, **Figure 2—figure supplement 1F**). A 3D invasion assay was also performed to check the ability of tumor spheres to invade a collagen matrix in YEATS2-knockdown and -overexpressed conditions. It was observed that a reduced number of cells invaded the collagen matrix when YEATS2 was downregulated. The opposite pattern was observed in both the cell lines when YEATS2 was overexpressed (**Figure 2F-G**, **Figure 2—figure supplement 1G-H**). All of the above observations led us to the conclusion that YEATS2 is one of the important epigenetic factors responsible for conferring head and neck cancer cells with invasive properties.

**Figure 2.**
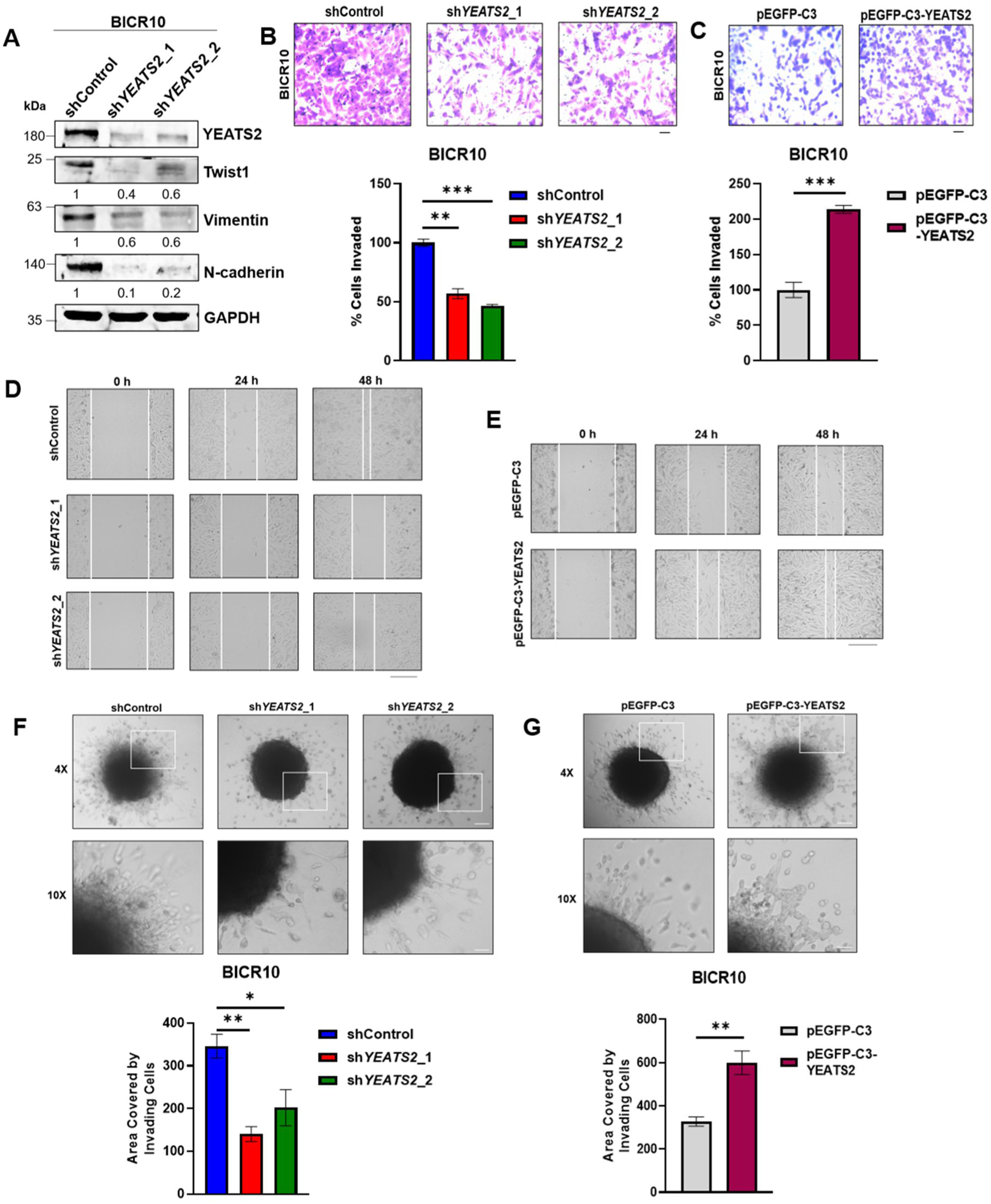
YEATS2 drives EMT in head and neck cancer cells. **(A)** Immunoblot showing the expression levels of various EMT factors upon YEATS2 knockdown in BICR10. **(B-C)** Results of invasion assay with quantification (below), after knockdown (B) or overexpression (C) of YEATS2 in BICR10 (Scale bar, 200 μm). **(D-E)** Wound healing assay performed after knockdown (D) or overexpression (E) of YEATS2 in BICR10 (Scale bar, 275 μm). **(F-G)** Results of 3D invasion assay showing change in invasive potential of BICR10 cells in collagen matrix after silencing (F) or overexpression (G) of YEATS2 (quantification shown below). (Scale bar: 4X, 200 μm; 10X, 50 μm) Error bars, mean ± SEM; two-tailed t test, ∗*p* < 0.05, ∗∗*p* < 0.01, ∗∗∗*p* < 0.001, n= 3 biological replicates.

### SP1 regulates the expression of YEATS2 in head and neck cancer

In order to investigate the molecular mechanism behind the upregulated expression of YEATS2 in HNC, YEATS2-promoter fragments of variable lengths were cloned upstream of Firefly luciferase reporter gene in the pGL3-Basic vector. Luciferase assay was performed in two different HNC cell lines, BICR10 and SCC9. A significant decrease in the luciferase activity was observed for shorter promoter-deletion construct (Luc-311) compared to the full-length construct (Luc-508) (**Figure 3A**, **Figure 3—figure supplement 1A)**. Transcription factor binding-site analysis was then performed with the sequence unique to Luc-508, the region potentially responsible for transcriptional regulation of YEATS2. This analysis revealed the presence of multiple consensus binding sites of SP1 and KLF5 transcription factors. Expression profiles of these transcription factors in the TCGA mRNA expression dataset for HNC showed that unlike *KLF5*, *SP1* was significantly upregulated in tumor vs. normal patient data (**Figure 3—figure supplement 1B-C**). Furthermore, shRNA-mediated silencing of SP1 led to decreased expression of YEATS2 in the HNC cell lines (**Figure 3B-C**, **Figure 3—figure supplement 1D-E**). Further, on performing SP1-ChIP assay, we found significant enrichment of SP1 on *YEATS2* promoter when compared with control IP in both the cell lines (**Figure 3D**, **Figure 3—figure supplement 1F**). Next, we also performed luciferase assay using Luc-508 construct in SP1-knockdown condition, and found a decreased luciferase activity as compared to control (**Figure 3E**). To further validate the direct involvement of SP1 in the transcriptional regulation of YEATS2, 3 of the 4 SP1 consensus binding sites in the full-length YEATS2 Luc-508 were mutated using site-directed mutagenesis. As expected, after performing luciferase assay, nearly 50% decrease in luciferase activity was observed for the mutant promoter construct, compared to its wild-type counterpart (**Figure 3F**, **Figure 3—figure supplement 1G**). We then looked into the status of invasion in sh*SP1* cells and found that the EMT-marker Twist1 was reduced in these cells. However, the ectopic expression of YEATS2 in sh*SP1* cells was able to rescue Twist1 expression partially (**Figure 3G**). We then performed invasion assay to check the effect of YEATS2 overexpression in the background of SP1-knockdown, and found that there was a decrease in invasion in sh*SP1* cells, which was rescued by overexpression of YEATS2 (**Figure 3H**, **Figure 3—figure supplement 1H**). These results indicate that SP1 is directly responsible for YEATS2 upregulation in HNC.

**Figure 3.**
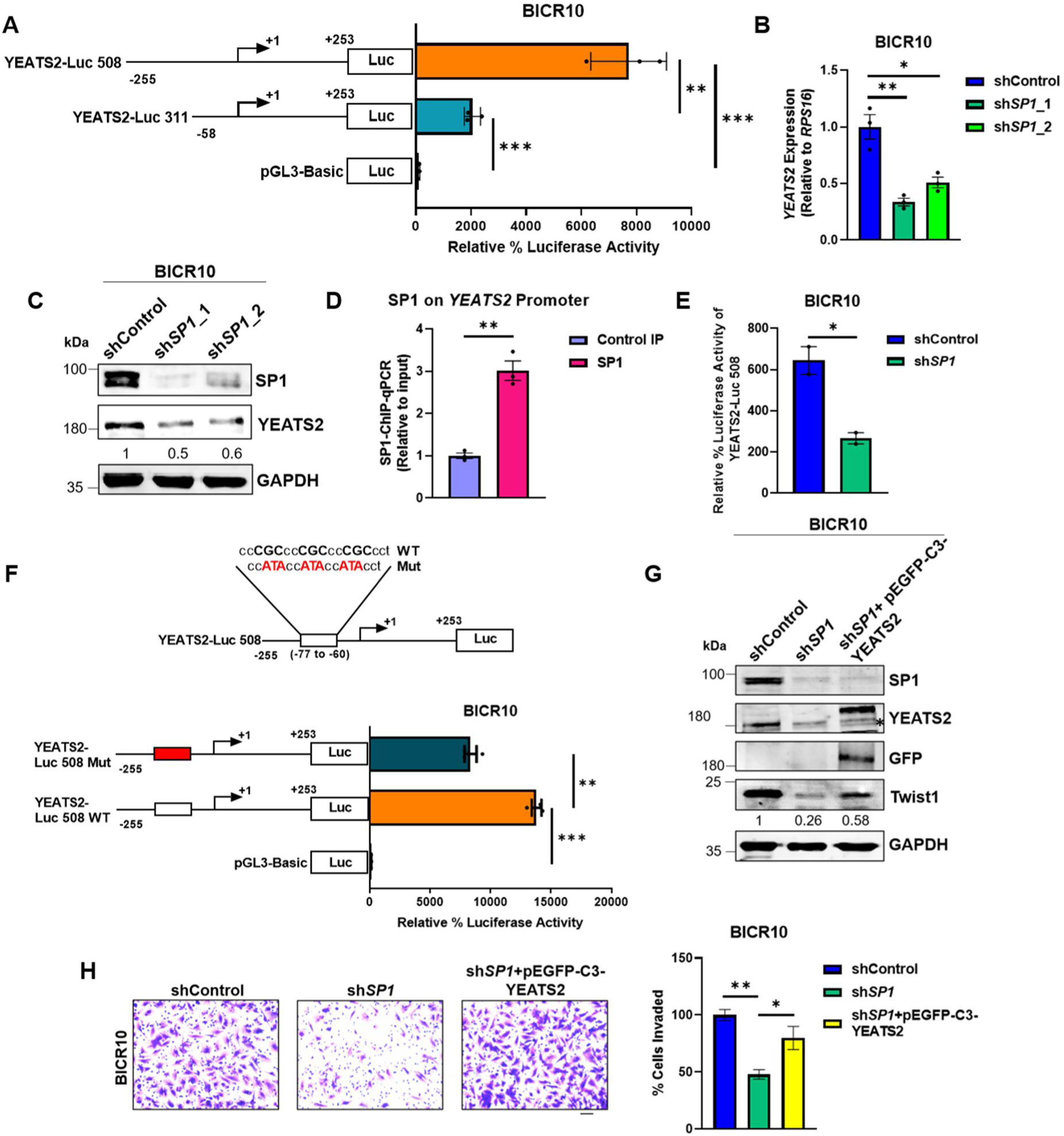
Regulation of EMT by YEATS2 is SP1-dependent in HNC. **(A)** Luciferase assay results showing the difference in relative luciferase activity of the two YEATS2 promoter deletion constructs in BICR10. **(B)** Plot showing decrease in mRNA expression of *YEATS2* on SP1-knockdown in BICR10 cells. **(C)** Immunoblot showing the reduced expression level of YEATS2 upon SP1-knockdown in BICR10. **(D)** Plot depicting SP1 binding on *YEATS2* promoter in SP1-ChIP-qPCR assay in BICR10. **(E)** Plot showing difference in relative luciferase activity of YEATS2 Luc-508 in shControl vs. sh*SP1* BICR10 cells. **(F)** Schematic showing mutation of SP1-binding site sequence in YEATS2 Luc-508 construct (above), and relative luciferase activity of wild-type (WT) vs. mutant (Mut) YEATS2 Luc-508 in BICR10 (below). **(G)** Immunoblot depicting the decreased Twist1 levels on SP1 knockdown and its subsequent rescue of expression upon YEATS2 overexpression in BICR10 (* indicates endogenous YEATS2 band). **(H)** Invasion assay images (with quantification on right) showing decrease and rescue of the percentage of invaded cells in sh*SP1* BICR10 cells, and sh*SP1* cells with YEATS2 overexpression, respectively. Scale bar, 200 μm. Error bars, mean ± SEM; two-tailed t test, ∗*p* < 0.05, ∗∗*p* < 0.01, ∗∗∗*p* < 0.001, n = 3 biological replicates.

### YEATS2 and GCDH regulate histone crotonylation in HNC

After establishing the role of YEATS2 as one of the important epigenetic factors involved in maintaining EMT-phenotype in HNC cells, we wanted to delve into the mechanism of YEATS2-mediated regulation of gene expression. Being a histone reader with a varying magnitude of affinity for multiple histone modifications, the function of YEATS2 could be potentially influenced by the metabolic state of the cell since metabolic intermediates can act as substrates for histone modifying proteins. Therefore, in order to explore the influence of cellular metabolism on the dynamics of histone modifications maintained by YEATS2, we wanted to look into the metabolic enzymes that are co-expressed with YEATS2 in HNC. For this, we performed gene set enrichment analysis (GSEA) with a comprehensive list of essential metabolic pathways ^18^ and found out that KEGG LYSINE DEGRADATION was the top-most pathway associated with high *YEATS2* expression in TCGA samples (**Figure 4A, Supplementary Table 3**). The saccharopine pathway (canonical pathway for lysine degradation) results in conversion of lysine to 2 molecules of acetyl-CoA, with several intermediates including crotonyl-CoA (**Figure 4B**) ^19^. Since it is known that YEATS2 has a higher affinity for binding crotonyl groups on histone (specifically on H3K27 residue) than other histone substrates ^9^, we hypothesized that the crotonyl-CoA producing enzyme glutaryl CoA dehydrogenase (GCDH), which is a part of lysine degradation pathway, would have increased expression in HNC. We found that *GCDH* was upregulated and had an inverse pattern of expression with the downstream enzyme *ECHS1* in TCGA cancer vs. normal tissue dataset (**Figure 4— figure supplement 1A-B**). Further, we found that the mRNA expression of *GCDH* was significantly upregulated in head and neck cancer samples relative to their matched normal tissues (**Figure 4C**). Moreover, correlation analysis performed between the expression levels of *GCDH* and *YEATS2* in TCGA data showed significant Spearman correlation coefficient value of 0.32 (**Figure 4—figure supplement 1C**). We then proceeded to check the levels of histone crotonylation (H3K27cr) in a set of nuclear protein lysates obtained from HNC tissues and found that the abundance of this histone mark was higher in tumor as compared to normal (**Figure 4D**, **Figure 4—figure supplement 1D**). Additionally, we performed IHC to evaluate the levels of ECHS1, GCDH, YEATS2, H3K27cr, and Twist1 in HNC tumor and their matched normal tissue sections. ECHS1 expression was notably reduced in tumor tissues, whereas GCDH, YEATS2, H3K27cr, and Twist1 exhibited increased staining intensity in tumor sections compared to the adjacent normal tissues (**Figure 4E**, **Figure 4—figure supplement 1E**). These findings indicate that the presence of YEATS2 and GCDH is possibly leading to an increase in the EMT phenotype (indicated by higher Twist1 expression) in cancer cells by promoting histone crotonylation. Furthermore, our observations suggest that an elevated GCDH/ECHS1 ratio in HNC tissues may be associated with increased levels of histone crotonylation. Since mitochondrial fraction of GCDH cannot influence histone crotonylation directly, we checked its subcellular localization in HNC tissues. Using immunohistochemistry, we detected the presence of nuclear GCDH in HNC tissues and observed that the regions that stained strongly for nuclear GCDH, also showed abundance in H3K27cr (**Figure 4—figure supplement 1F**). Conversely, the weakly stained regions for nuclear GCDH were less abundant in H3K27cr levels. This established an association between the presence of H3K27cr marks and the expression of GCDH in HNC. Further, on checking the levels of H3K27cr on YEATS2-and GCDH-knockdown using immunoblotting and immunofluorescence, we found that the cells silenced for either of the two genes caused a decrease in H3K27cr levels (**Figure 4F-I**, **Figure 4—figure supplement 1G-H**). On the other hand, an increase in H3K27cr signal was observed when YEATS2 was overexpressed ectopically in BICR10 cells (**Figure 4—figure supplement 1I**). These findings indicate a coordinated regulation of histone crotonylation in HNC cells by the epigenetic factor YEATS2 and the metabolic enzyme GCDH.

**Figure 4.**
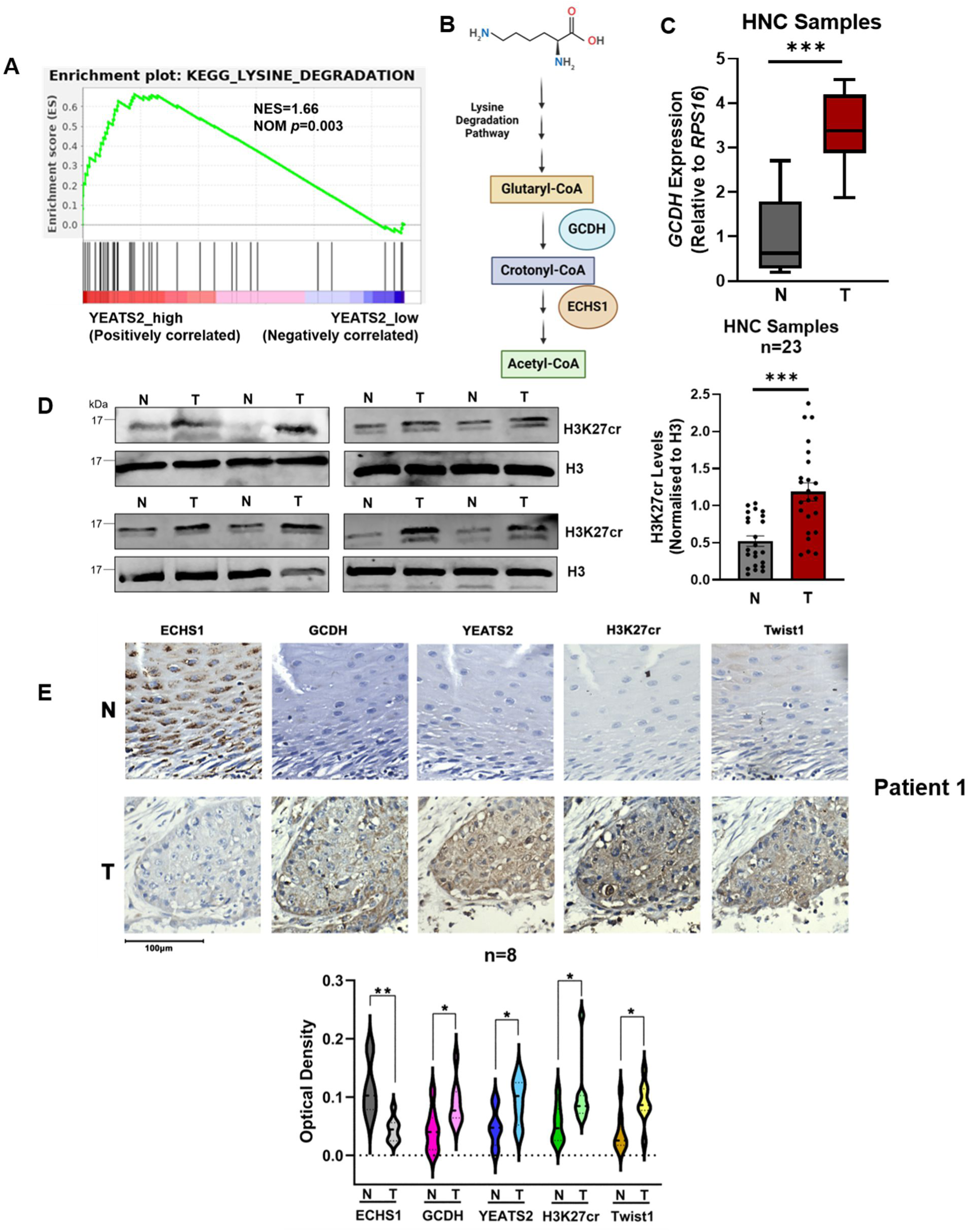
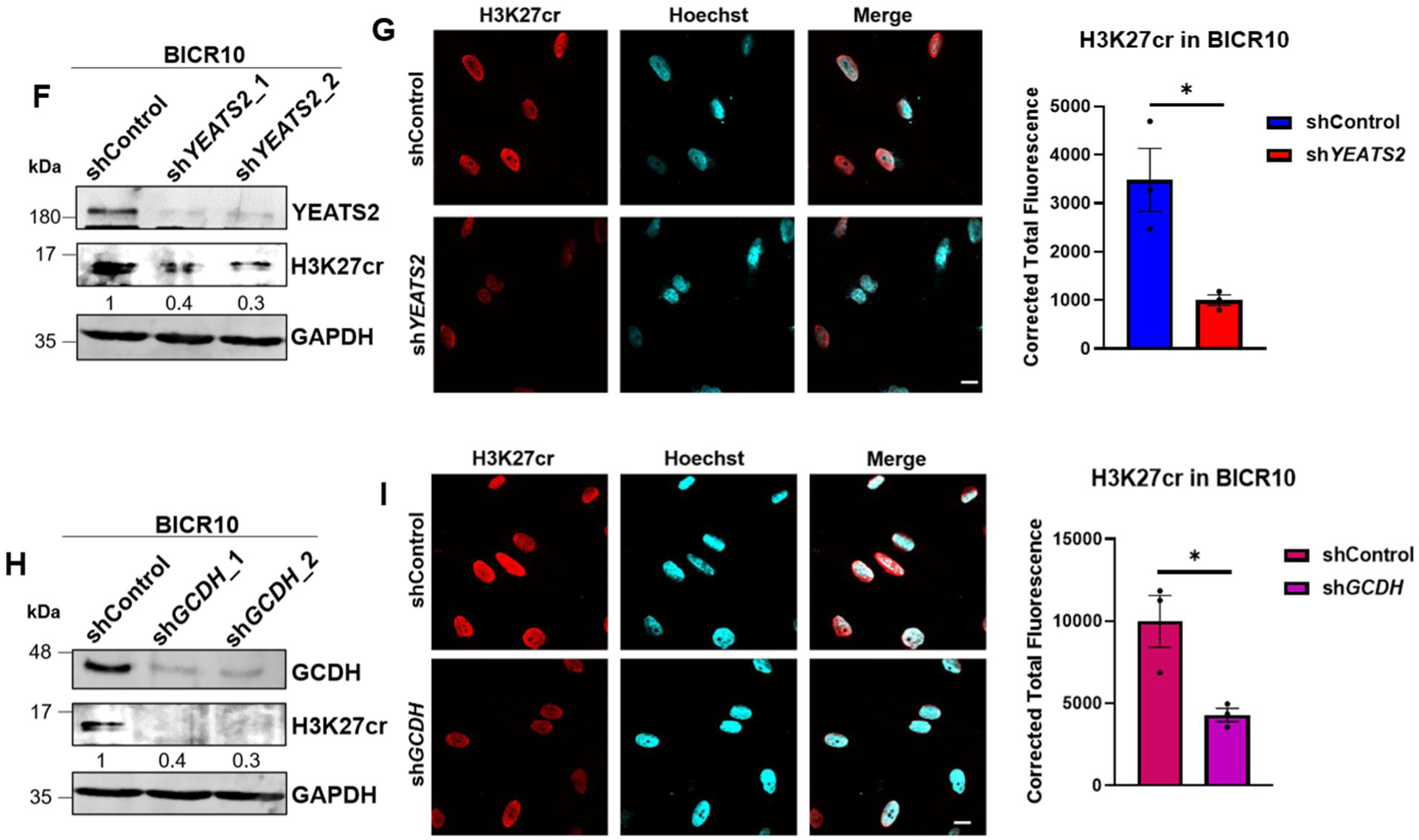
YEATS2 and GCDH regulate histone crotonylation in HNC. **(A)** GSEA plot showing enrichment of KEGG LYSINE_DEGRADATION_PATHWAY in TCGA samples stratified as YEATS2_high as compared to YEATS2_low samples (NES-net enrichment score, NOM p-nominal p-value). **(B)** Schematic depicting the canonical lysine degradation pathway highlighting the step that leads to crotonyl-CoA production via GCDH. **(C)** RT-qPCR result showing enhanced mRNA expression of *GCDH* in tumor vs. normal samples (n=8). **(D)** Immunoblot showing enhanced levels of H3K27cr (quantification on right) in nuclear lysates extracted from HNC tumor vs. normal samples (n=23). **(E)** Representative IHC images (quantification below) showing the levels of ECHS1, GCDH, YEATS2, H3K27cr and Twist1 in HNC normal vs. tumor tissue samples (n=8) (Scale bar, 100 μm). **(F and H)** Immunoblot depicting the decrease in H3K27cr levels on (F) YEATS2- and (H) GCDH-knockdown in BICR10 cells. **(G and I)** Immunoflorescence images depicting the decrease in H3K27cr levels on (G) YEATS2- and (I) GCDH-knockdown in BICR10 cells (Scale bar, 10 μm). Error bars, mean ± SEM; two-tailed t test, ∗*p* < 0.05, ∗∗*p* < 0.01, ∗∗∗*p* < 0.001. n = 3 biological replicates.

### YEATS2 regulates expression of EMT-associated gene SPARC

In order to check the effect of YEATS2 on global gene expression, we performed YEATS2 ChIP-seq to obtain genome-wide binding sites of this protein (**Figure 5— figure supplement 1A, Supplementary Table 4**). We then performed integration of those genes having YEATS2 ChIP-seq peaks on their promoters with the genes that were downregulated in sh*YEATS2* vs. shControl RNA-seq data, and the genes included in the hallmark EMT gene signature. We found that *SPARC* was the common gene present in all 3 sets (**Figure 5A-B**, **Figure 5—figure supplement 1B**). The mRNA expression level of *SPARC* in TCGA dataset was also found to be higher in tumor as compared to normal (**Figure 5—figure supplement 1C**). Therefore, we chose to further investigate *SPARC* as a direct gene target of YEATS2 in mediating EMT in HNC. Secreted protein acidic and rich in cysteine (SPARC) is a secreted glycoprotein known to be a part of extracellular matrix (ECM). It interacts with various components of ECM and is known to play a role in processes like tissue remodeling and angiogenesis ^20,21^. By performing YEATS2 ChIP-qPCR in YEATS2-downregulated cells, we found that the binding of YEATS2 on *SPARC* promoter decreased on its knockdown in BICR10 as well as SCC9 (**Figure 5C**, **Figure 5—figure supplement 1D**). We then checked the expression of *SPARC* at the RNA level on YEATS2-downregulation. *SPARC* expression was decreased in YEATS2-knockdown conditions in both the cell lines (**Figure 5D**, **Figure 5—figure supplement 1E**). Since SPARC is a secretory protein, we investigated its level in conditioned media obtained from shControl and sh*YEATS2* BICR10 cells. We found a decrease in the secreted pool of SPARC on YEATS2-knockdown (**Figure 5E**). We also observed a decrease in the cellular levels of SPARC protein in SCC9 (**Figure 5—figure supplement 1F**). Since the binding of a histone reader protein to a gene promoter cannot solely lead to its increased expression, we explored the possibility of the involvement of a writer protein in collaborating with YEATS2 to maintain higher levels of SPARC in HNC. For this, we checked the expression of p300, a known writer of H3K27cr mark ^22^ in humans, in TCGA dataset. We found that p300 (*EP300* gene) had increased expression in tumor vs. normal HNC dataset (**Figure 5—figure supplement 1G**). On performing p300-ChIP assay, it was found that its binding was reduced in YEATS2-silenced BICR10 cells when compared to the shControl cells, indicating the potential role of p300 in leading to increased expression of SPARC in HNC cells (**Figure 5F**). To validate our claim, we performed immunoblotting after p300 knockdown and found that the expression of SPARC in supernatant derived from sh*EP300* BICR10 cells was reduced as compared to shControl cells (**Figure 5G**). Further, we performed p300 pulldown in a co-immunoprecipitation experiment to check if YEATS2 and p300 function in a protein complex. We discovered the presence of YEATS2 in the p300-immunoprecipitate sample, clearly indicating that YEATS2 and p300 function together as a complex to drive the expression of SPARC (**Figure 5H**). This set of results indicates a sophisticated epigenetic mechanism promoting increased transcription of the EMT-related gene SPARC in HNC. Finally, we wanted to check the effect of SPARC on invasion. It was observed that the decrease in invasion seen on YEATS2-downregulation was rescued by SPARC overexpression (**Figure 5I-J**). This indicates that SPARC is a direct target of YEATS2 and acts as a downstream effector of YEATS2 in maintaining invasive phenotype in HNC cells.

**Figure 5.**
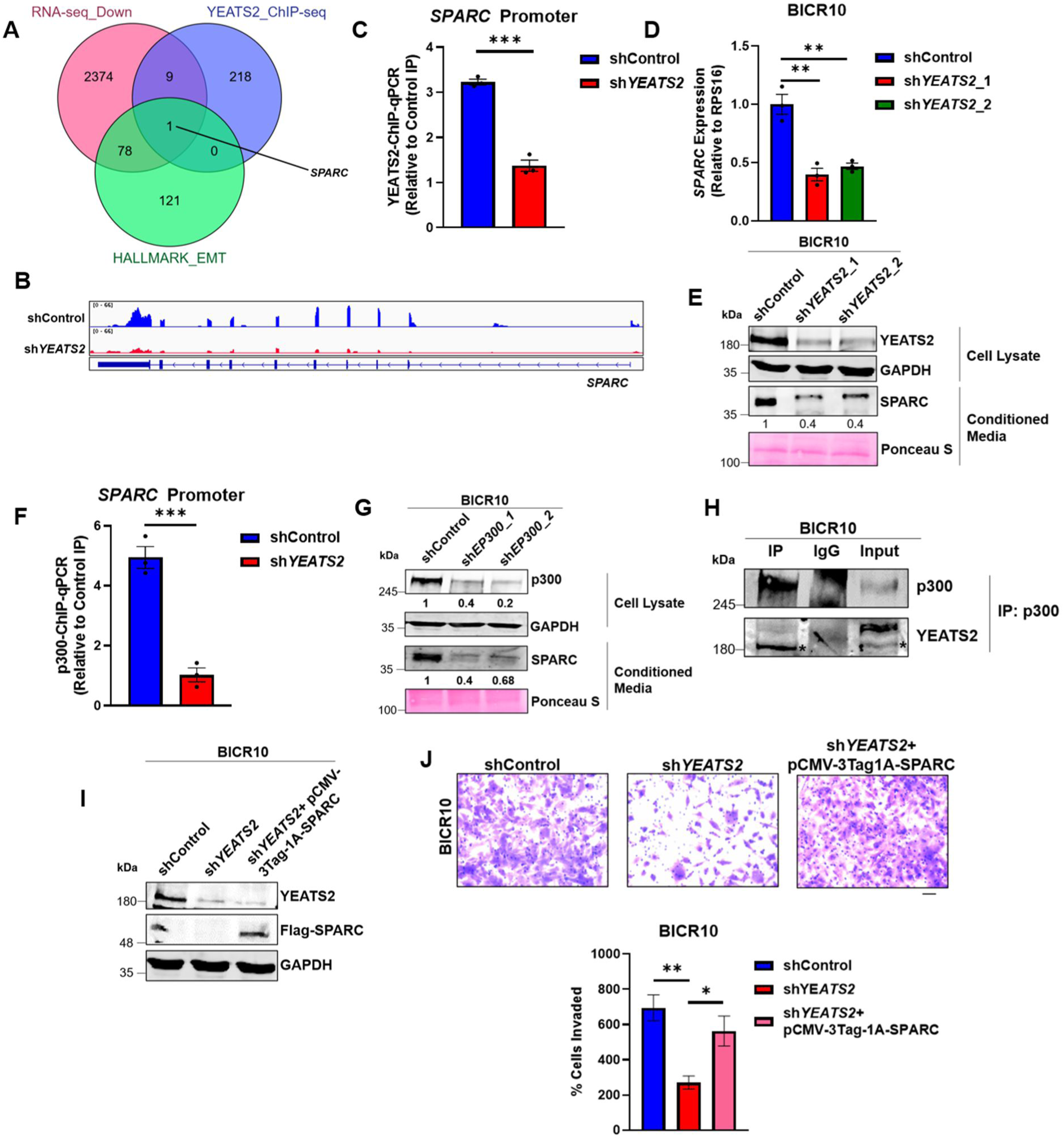
YEATS2 regulates expression of EMT-related SPARC in HNC. **(A)** Venn diagram showing *SPARC* gene obtained after integration of RNA-seq data (genes downregulated in shControl vs. sh*YEATS2*), YEATS2 ChIP-seq data and, hallmark EPITHELIAL_MESENCHYMAL_TRANSITION gene signature. **(B)** Integrative genome viewer (IGV) plot showing decrease in SPARC expression in shControl vs. sh*YEATS2* RNA-seq data. **(C)** YEATS2-ChIP-qPCR results showing decreased binding of YEATS2 on *SPARC* promoter in sh*YEATS2* BICR10 cells. **(D)** RT-qPCR results showing decreased expression of *SPARC* on YEATS2-knockdown in BICR10. **(E)** Immunoblot showing decreased expression of SPARC in conditioned media derived from shControl and sh*YEATS2* BICR10 cells. **(F)** p300-ChIP-qPCR results showing decreased binding of p300 on *SPARC* promoter in sh*YEATS2* BICR10 cells. **(G)** Immunoblot showing decreased SPARC levels on p300 knockdown in conditioned media from shControl and sh*EP300* BICR10 cells. **(H)** Immunoblot showing co-immunoprecipitation of YEATS2 by p300 in YEATS2-overexpressed BICR10 cells (endogenous YEATS2 band is highlighted by *). **(I-J)** Immunoblot depicting the overexpression of Flag-tagged SPARC in sh*YEATS2* BICR10 cells (I), and Invasion assay images (J) (with quantification below) showing decrease and rescue of the percentage of invaded cells in shControl vs. sh*SP1* BICR10 cells, and sh*SP1* cells with YEATS2 overexpression, respectively. Scale bar, 200 μm. ∗∗*p* < 0.01, ∗∗∗*p* < 0.001, n = 3 biological replicates.

### Maintenance of H3K27cr marks is dependent on YEATS2-mediated recruitment of p300 on *SPARC* promoter

To investigate the mechanism of YEATS2-mediated gene regulation we checked the status of histone modifications, both acetylation and crotonylation at the H3K27 residue. We performed ChIP-qPCR assay to check the abundance of both the marks at the *SPARC* promoter and found that only H3K27cr mark showed significant downregulation on YEATS2 knockdown (**Figure 6A-B**, **Figure 6—figure supplement 1A-B)**. This indicated that SPARC expression is potentially controlled by a non-acetyl histone mark i.e., crotonylation. Further, to validate the role of YEATS2 in the transcription of *SPARC* gene, we performed RNA Pol II ChIP-qPCR assay and observed a decrease in Pol II occupancy on *SPARC* promoter after YEATS2 knockdown (**Figure 6C**). Since the substrate used in addition of crotonylation marks on histones is crotonyl-CoA, we supplemented the cells with its precursor molecule sodium crotonate ^23^, and saw that there was an increase in H3K27cr levels on *SPARC* promoter (**Figure 6D**, **Figure 6—figure supplement 1C**). We also observed an increase in the expression of SPARC at mRNA and protein levels in both the cell lines (**Figure 6E-F**, **Figure 6—figure supplement 1D-E**). We then checked the effect of crotonate on cancer cell invasiveness and observed that cells treated with crotonate invaded in greater numbers in Matrigel than untreated cells in both the HNC cell lines (**Figure 6G**, **Figure 6—figure supplement 1F**). Finally, we checked whether crotonate treatment can rescue the effect of YEATS2 downregulation on SPARC expression. We did not observe significant rescue either in H3K27cr levels on *SPARC* or in SPARC protein expression, when YEATS2-downregulated cells treated with crotonate (**Figure 6H-I**). With these observations, we can conclude that H3K27cr-mediated increase in SPARC expression is subject to the availability of YEATS2 in HNC cells.

**Figure 6.**
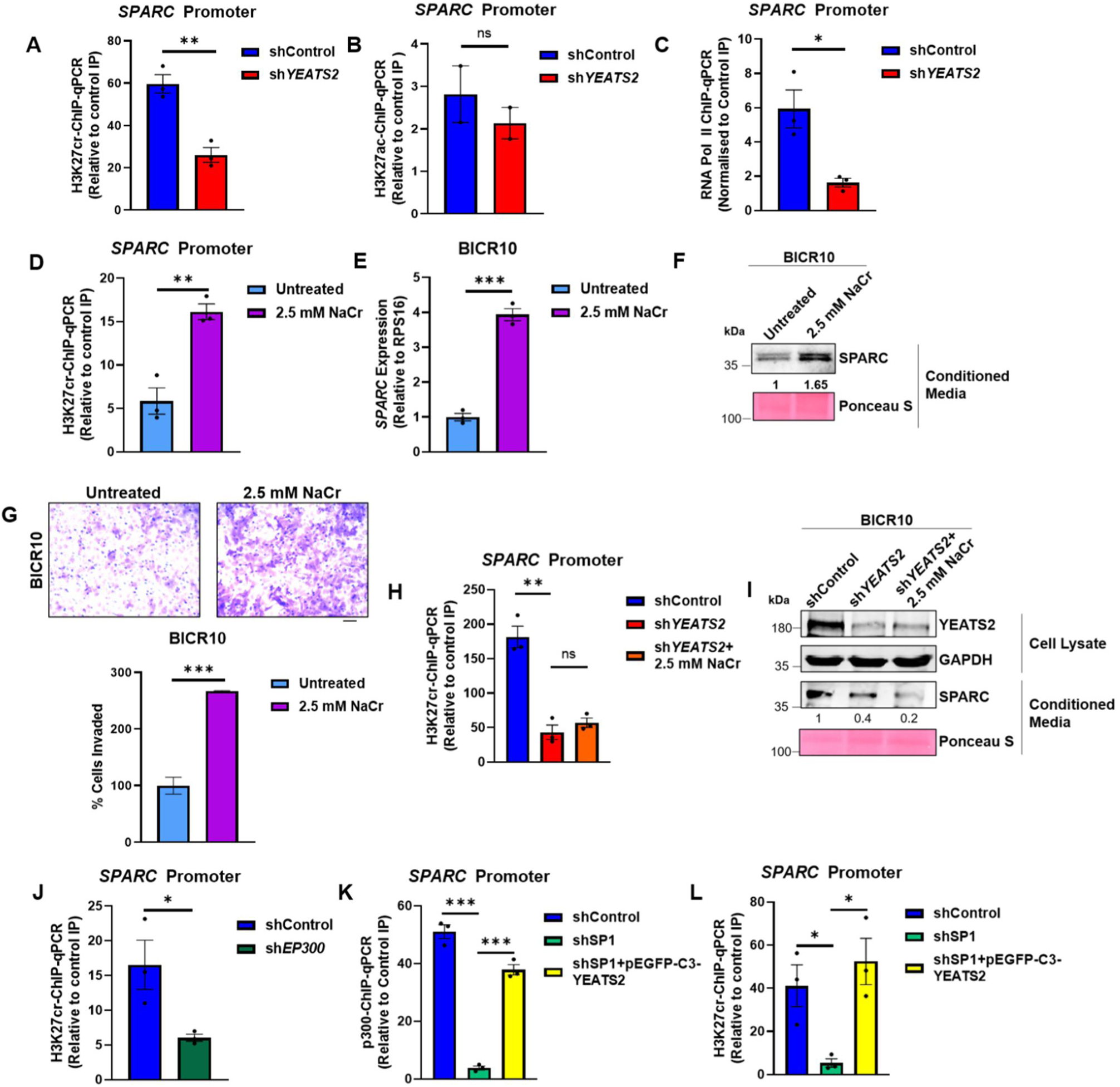
Maintenance of H3K27cr marks is dependent on YEATS2-mediated recruitment of p300 on SPARC promoter. **(A)** H3K27cr-ChIP-qPCR results showing decrease in H3K27cr enrichment on *SPARC* promoter on YEATS2 knockdown. **(B)** H3K27ac-ChIP-qPCR results showing non-significant change in H3K27ac enrichment on *SPARC* promoter on YEATS2 knockdown. **(C)** RNA Pol II ChIP-qPCR showing decrease in Pol II occupancy on SPARC after YEATS2 downregulation in BICR10. **(D)** H3K27cr-ChIP-qPCR results showing increase in H3K27cr enrichment on *SPARC* promoter on treating BICR10 cells with 2.5 mM sodium crotonate (NaCr). **(E-F)** RT-qPCR (E) and immunoblot (F) showing enhanced SPARC expression in untreated BICR10 cells vs. BICR10 cells treated with 2.5 mM NaCr. **(G)** Invasion assay images (with quantification below) showing increased invasion on treating BICR10 cells with 2.5 mM NaCr (Scale bar, 200 μm). **(H)** H3K27cr-ChIP-qPCR data depicting the lack of significant difference in H3K27cr levels between sh*YEATS2* vs. sh*YEATS2*+ 2.5 mM NaCr cells. **(I)** Immunoblot showing inability of NaCr treatment to rescue SPARC expression in YEATS2-knockdown BICR10 cells. **(J)** H3K27cr ChIP-qPCR showing decrease in H3K27cr levels in BICR10 shControl vs. sh*EP300* cells. **(K-L)** p300 (K) and H3K27cr ChIP-qPCR (L) showing decrease and subsequent rescue in p300 binding and H3K27cr enrichment on SP1 knockdown and YEATS2 overexpression, respectively. Error bars, mean ± SEM; two-tailed t test, ns-non-significant, ∗*p* < 0.05, ∗∗*p* < 0.01, ∗∗∗*p* < 0.001.

Since we have previously established the role of p300 in the transcriptional regulation of SPARC, we now wanted to investigate the exact mechanism involved behind this regulation. For this, we performed p300 knockdown followed by H3K27cr-ChIP-qPCR and found a significant decrease in H3K27cr levels on *SPARC* promoter in sh*EP300* BICR10 cells (**Figure 6J**). This indicates that p300 regulates SPARC expression by functioning as a crotonyltransferase. Further, we wanted to check whether the binding and subsequent activity of p300 is dependent on the presence of YEATS2 on *SPARC* promoter. As shown in Figure 3, YEATS2 expression in HNC cells is dependent on SP1. To examine the dependency of p300 activity on YEATS2, we chose to knockdown SP1 followed by YEATS2 overexpression. As expected, on performing p300-ChIP-qPCR we saw a decrease in p300 occupancy on *SPARC* in SP1 knockdown cells, which was rescued by overexpressing YEATS2 in sh*SP1* cells (**Figure 6K**). A decrease in SP1 levels would have led to abrogation of YEATS2, which in turn reduced p300 recruitment at the *SPARC* promoter. We observed the same pattern in the H3K27cr levels—decreased enrichment of H3K27cr in sh*SP1* cells and its subsequent rescue on YEATS2 overexpression (**Figure 6L**). In conclusion, the expression of EMT-related gene SPARC is regulated by promoter H3K27cr levels added by crotonyltransferase p300, whose recruitment is dependent on the histone reader YEATS2.

### GCDH expression is SP1-dependent and it regulates H3K27cr-mediated SPARC expression with YEATS2 synergistically

As previously shown, both YEATS2 and GCDH regulate the levels of histone crotonylation in HNC. So, we hypothesized that the presence of GCDH could be essential for production of crotonyl-CoA, which could be added to histones by histone modification machinery consisting of YEATS2 and p300. On performing knockdown of GCDH, we observed a decrease in SPARC at protein and mRNA levels (**Figure 7A-B**, **Figure 7—figure supplement 1A**). Also, on performing H3K27cr-ChIP assay with GCDH-downregulated cells, we found that H3K27cr marks on *SPARC* promoter reduced relative to control cells (**Figure 7C**). We then sought to explore the possibility of localized histone crotonylation performed by a multi-protein complex consisting of YEATS2 and GCDH. For this, we performed co-immunoprecipitation; however, we did not observe an interaction between GCDH and YEATS2 (**Figure 7—figure supplement 1B**), ruling out the possibility of the localized production of crotonyl-CoA at a specific gene locus ^24^. We then checked whether both the genes are regulated by the same transcription factor i.e., SP1. TCGA gene expression dataset showed a significant correlation between the expression levels of *SP1* and *GCDH*, hinting at the possibility of SP1 transcriptionally regulating GCDH in HNC (**Figure 7—figure supplement 1C**). Thus, we performed SP1-knockdown and found a decrease in GCDH levels in both HNC cell lines (**Figure 7D**, **Figure 7—figure supplement 1D**). On performing SP1-ChIP we found a significant occupancy of SP1 on *GCDH* promoter in BICR10 and SCC9 (**Figure 7E**, **Figure 7—figure supplement 1E**). These results suggest that SP1 is responsible for high expression of both GCDH and YEATS2 in HNC. As we have already established that YEATS2 abrogation leads to decrease in the invasiveness of HNC cells, we now wanted to check the effect of GCDH downregulation on HNC cell invasion. We performed Matrigel invasion assay with shControl and sh*GCDH* BICR10 cells and found a significant decrease in the invasive capacity of sh*GCDH* cells (**Figure 7F**). Conversely, the number of invading cells showed an increase when GCDH was overexpressed in BICR10 cells (**Figure 7—figure supplement 1F**), indicating that like YEATS2, GCDH is also essential for imparting invasive properties to HNC cells.

**Figure 7.**
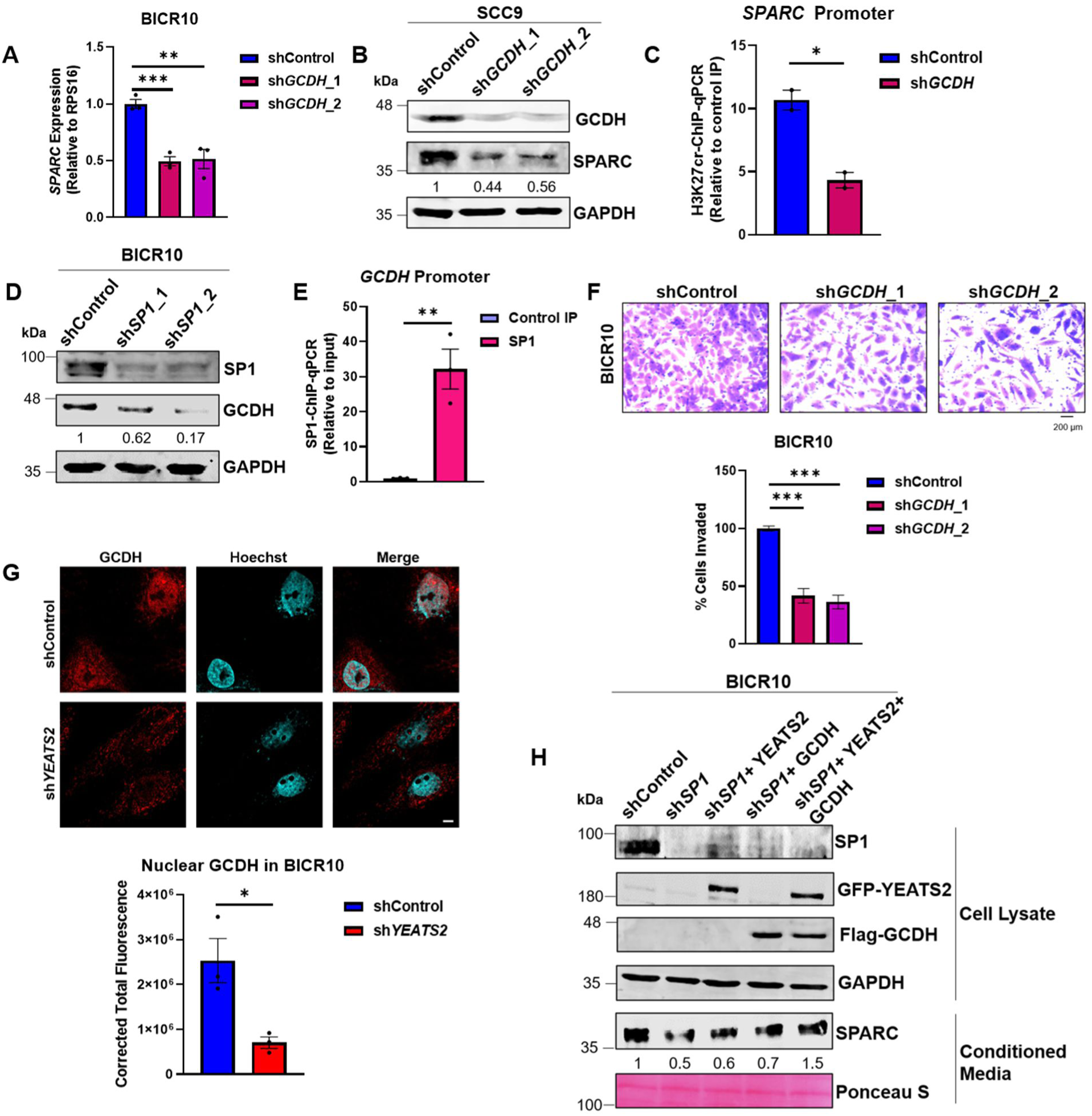
GCDH expression is SP1-dependent and regulates H3K27cr-mediated SPARC expression with YEATS2 synergistically. **(A)** RT-qPCR results showing decrease in *SPARC* expression on GCDH-knockdown in BICR10 cells. **(B)** Immunoblot showing the reduced expression of SPARC upon GCDH-knockdown in SCC9 cells. **(C)** Decrease in H3K27cr levels on *SPARC* promoter in sh*GCDH* BICR10 cells. **(D)** Immunoblot showing the reduced expression of GCDH on SP1-knockdown in BICR10 cells. **(E)** Plot showing SP1 binding on *GCDH* promoter in SP1-ChIP assay in BICR10. **(F)** Invasion assay images (quantification shown below) showing decrease in invasion of BICR10 cells on GCDH knockdown (Scale bar, 200 μm). **(G)** IF images (quantification shown below) showing reduced nuclear localization of GCDH in sh*YEATS2* BICR10 cells. Scale bar, 5 μm. **(H)** Immunoblot depicting decreased SPARC expression on SP1 knockdown and its subsequent rescue upon dual YEATS2 and GCDH overexpression in sh*SP1* BICR10 cells. Error bars, mean ± SEM; two-tailed t test, ∗*p* < 0.05, ∗∗*p* < 0.01, ∗∗∗*p* < 0.001, n = 3 biological replicates.

The canonical location of GCDH in lysine degradation pathway is mitochondria. However, having detected the presence of nuclear pool of GCDH in HNC tissues (**Figure 4—figure supplement 1F**), we wanted to investigate the role of YEATS2 in affecting the subcellular localization of GCDH in HNC cells. We detected the presence of GCDH in nucleus in BICR10 using IF, and observed that the levels of nuclear GCDH diminished in sh*YEATS2* cells as compared to shControl cells (**Figure 7G**). This indicates that YEATS2 might be involved in regulating the transport of GCDH inside nucleus indirectly by an unknown mechanism. In order to further investigate this epigenome-metabolism interplay, we checked the role of YEATS2 and GCDH in regulating SPARC expression in the absence of SP1. The dual overexpression of YEATS2 and GCDH in SP1-silenced cells rescued the SPARC levels to a greater degree; whereas, YEATS2 and GCDH when overexpressed alone, could only partially rescue SPARC expression (**Figure 7H**). Hereby, we establish that one of the axes involved in regulation of EMT in HNC is dependent on the synergistic maintenance of H3K27cr marks on *SPARC* gene by YEATS2 and GCDH.

### YEATS2 regulates gene expression by maintaining H3K27cr levels globally

We have previously shown that the expression of a large number of genes are affected by YEATS2 downregulation (**Figure 1H**). For establishing a direct relationship between YEATS2-mediated histone crotonylation and gene expression, we performed ChIP-sequencing to check global H3K27cr levels in shControl and sh*YEATS2* BICR10 cells. We found that downregulation of YEATS2 led to a significant change in H3K27cr enrichment at 1,583 locations in the genome, out of which 1,533 locations showed significant decrease in H3K27cr enrichment (**Figure 8A**). Further, 1,166 differentially lost sites were found to be present at the promoter regions of genes. As expected, overrepresentation analysis using these set of genes revealed EPITHELIAL MESENCHYMAL TRANSITION as one of the enriched hallmark gene sets (**Figure 8B**). We then overlapped the genes having reduced H3K27cr levels with the genes that showed downregulation in shControl vs. sh*YEATS2* RNA-seq. A total of 147 genes were found to be common in both the sets (**Figure 8C, Supplementary Table 5**). Among these 147 genes, some examples of EMT-associated genes were *ERG* ^25^ and *CREB3L2* ^26^ (**Figure 8D-E**). Overall, our results suggest that YEATS2 regulates histone crotonylation throughout the genome resulting in the transcriptional activation of a large number of genes, many of which are related to the process of EMT.

**Figure 8.**
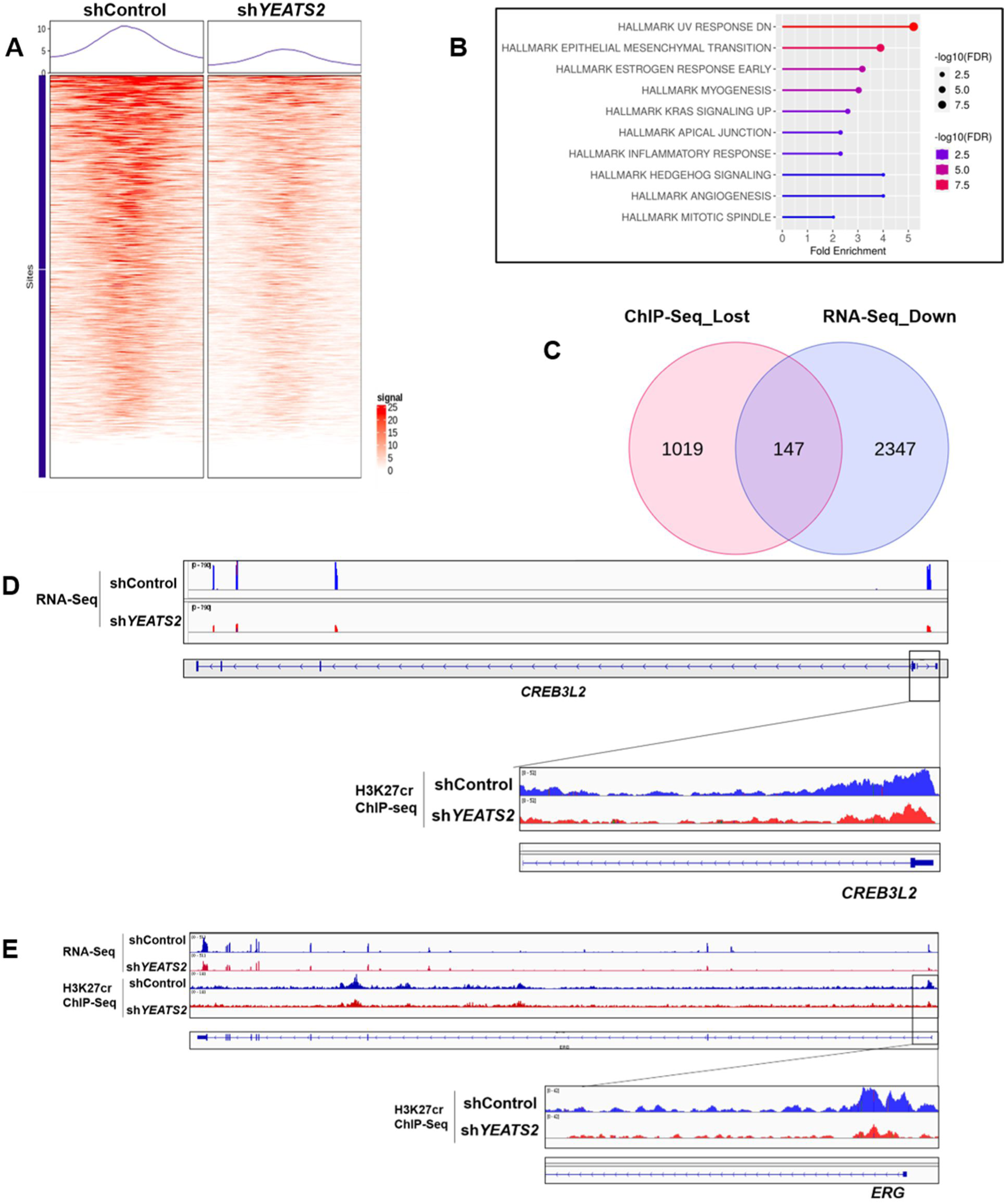
YEATS2 regulates gene expression by maintaining H3K27cr levels globally. **(A)** Differential ChIP-seq profile of H3K27cr in shControl vs. sh*YEATS2*. **(B)** Overrepresentation analysis showing pathways enriched among genes with decreased H3K27cr enrichment in H3K27cr ChIP-seq data. **(C)** Overlap of genes with reduced H3K27cr on their promoter and downregulated genes in RNA-seq. **(D-E)** IGV plot showing representative examples of genes (CREB3L2 and ERG) common in (C).

## Discussion

In our study, we identified YEATS2 as an important epigenetic regulator that plays an oncogenic role in head and neck tumor progression. We have shown that the histone reader regulates expression of important EMT-related genes. The mechanism employed by YEATS2 for gene regulation was also investigated. The YEATS domain confers unique properties to this epigenetic factor, making it suitable for binding with modifications other than acetylation ^27^. As stated earlier, YEATS2 has a binding affinity for a variety of chemical groups. Among all histone modifications screened, YEATS2 had the highest affinity for histone crotonylation, specifically at the histone H3 lysine 27 position ^9^Click or tap here to enter text.. There have been conflicting reports about the role of crotonylation at the H3K27 position. YEATS domain-containing GAS41 was found to recruit SIN3A-HDAC1 complex in a H3K27cr-dependent manner at the promoters of tumor suppressor genes, thereby leading to their repression in colorectal cancer ^28^. Conversely, another study in colorectal cancer showed that high H3K27cr at the promoter of *ETS1* gene led to an increase in its expression and promoted metastasis ^10^. In our findings, we have corroborated the latter study by showing the association of H3K27cr with the activation of an EMT-associated gene.

With the discovery of a plethora of novel histone modifications, researchers have been facing the challenge to understand the role of these marks in influencing gene expression. A group of such histone marks, known as the non-acetyl acylation marks, consist of a wide variety of chemical moieties which differ in structure and chemical properties ^29,30^. Due to the differences amongst these histone marks, it is obvious to assume that the proteins interacting with such histone modifications would not be the same as those interacting with histone acetylation or methylation. YEATS domain is one of the domains that has affinity for non-acetyl histone acylation marks. There are 4 members in the family of YEATS-domain containing proteins, AF9, YEATS2, ENL and, GAS41 ^27^. By using multiple high-throughput next generation sequencing techniques, we have strengthened the evidence for the role of YEATS2 in upregulating the expression of EMT-associated gene targets by maintaining a non-acetyl histone mark on their promoters. Global abrogation of pro-EMT genes was seen in RNA-seq data of YEATS2-silenced cells, which was underlaid by a simultaneous decrease seen in H3K27cr levels observed in H3K27cr ChIP-seq. This established a direct relationship between tumorigenic gene expression and histone crotonylation in cancer. Furthermore, in order to thoroughly investigate this link, we have integrated different sets of NGS data and found that invasion-associated gene *SPARC* gene is a direct target of H3K27cr-mediated regulation by YEATS2, and is possibly one of the many ways in which the EMT-promoting effect of YEATS2 is relayed downstream. As a part of the tumor microenvironment, SPARC mediates ECM-tumor cell communication. Several reports have suggested the role of SPARC in contributing towards a pro-invasive phenotype in cells multiple cancer types ^31,32^. Presence of SPARC in tumor stroma was reported to be responsible for recruitment of immunosuppressive myeloid cells in high grade breast carcinomas ^33^. In this study, we observed that the SPARC was capable of rescuing the decrease in invasive properties of YEATS2-silenced cells, making it the functional downstream effector of YEATS2.

We have discovered direct interaction between YEATS2 and epigenetic writer p300 in our study. Research has demonstrated diversity among histone reader families, each exhibiting preferences for specific histone marks. However, considerable overlap exists among histone marks deposited by the same histone writer. Notably, we found that one of the most promiscuous acyltransferases, p300, can deposit H3K27cr marks at the SPARC gene promoter in a YEATS2-dependent manner. Using ChIP assays and rescue experiments we showed that p300 binds to SPARC promoter and its binding is reduced in the absence of YEATS2. Recent studies have highlighted the importance of p300-mediated crotonylation at H3K27 and other histone residues in the crucial process of early embryonic development. Depletion of p300 led to reduced histone crotonylation resulting in dysregulated gene expression in developing embryos causing them to be arrested at the 8-cell stage ^34^. Our study adds to the growing body of research underscoring the importance of YEATS2-and p300-mediated histone crotonylation in both physiological and pathological processes.

Our study demonstrates the cooperation between an epigenetic factor and a metabolic enzyme working together to mediate EMT-promoting gene expression. This crosstalk has not been reported in head and neck cancer previously. Since epigenetic changes and aberrant metabolism both are major hallmarks of cancer ^8,35^, it is important to investigate the metabolism-epigenetics axis in order to yield new therapeutic targets. Increasing evidences suggest that the state of chromatin in a cell is a reflection of the metabolic enzymes functioning inside the nucleus. Through gene set enrichment analysis, we found that there was a significant correlation between YEATS2 and the lysine degradation pathway in HNC. This led to the observation that one of the enzymes crucial for the production of crotonyl-CoA, GCDH is overexpressed in HNC. Previous reports have suggested that metabolic enzymes could affect epigenomic changes by functioning closely with epigenetic factors. Studies showing the presence of multi-subunit complexes consisting of a metabolite-producing enzyme, DNA-binding protein and, a histone writer are a testament to the fact that metabolism-epigenome collaboration is more direct than previously thought ^36^. Although we did not observe direct association of GCDH and YEATS2, we did establish that the nuclear presence of GCDH causes an increase in histone crotonylation in HNC. As per the evidence provided by us, it could be proposed that the surge in GCDH in YEATS2-high cancer cells affects the chromatin by increasing the pool of crotonyl-CoA inside nucleus, which acts as a substrate for histone modifying enzymes. Since YEATS2 works by recruiting p300 via recognition of histone marks, maintenance of crotonylation at promoter regions could be dependent on sites with a constitutive basal level of H3K27cr or other marks, as a histone reader cannot recognize a genomic locus devoid of pre-existing histone marks. Such constitutive sites would be used by YEATS2 as an anchor to recruit p300 and subsequently induce histone crotonylation on nearby “inducible” sites. Further investigation is needed to precisely locate and distinguish between constitutive and inducible sites of histone crotonylation. Moreover, we have shown that SP1 transcriptionally regulates both YEATS2 and GCDH. This hints towards a possibility of a direct axis initiated by an unknown upstream factor working in order to modify the chromatin in such a manner that it benefits the cancer cell by causing an increase in the tumorigenic properties of the cell.

In conclusion, we have presented a sophisticated interplay between YEATS2 and GCDH, where the upregulation of both factors contributes towards the enhancement of EMT in head and neck cancer by changes in the global EMT-favoring gene expression, mediated through promoter histone crotonylation (**Figure 9**). Our findings provide new insights into the metabolic–epigenetic crosstalk driving EMT and identify the YEATS2–GCDH–SPARC axis as a potential therapeutic target in HNC.

**Figure 9.**
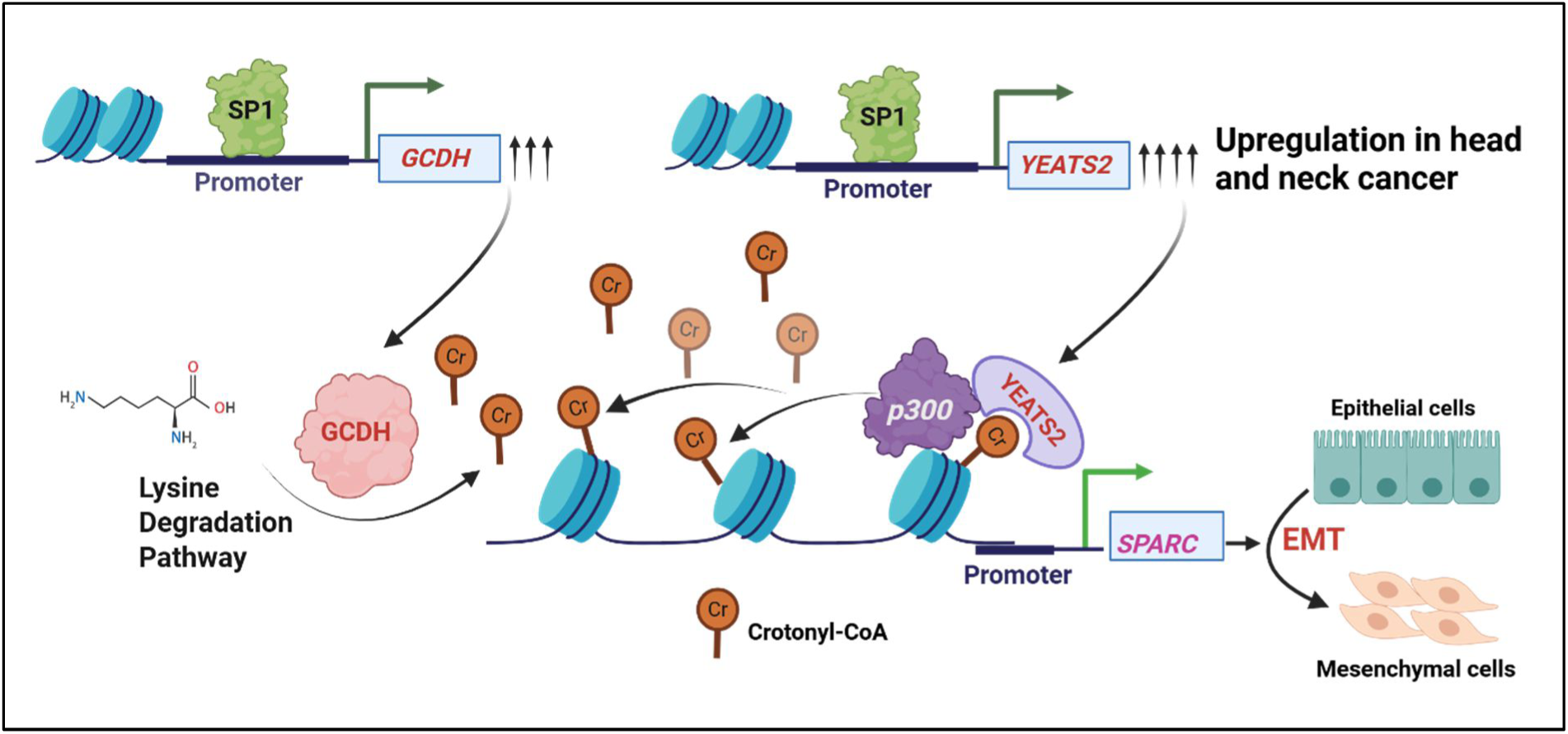
Schematic showing YEATS2- and GCDH-mediated regulation of EMT through H3K27cr-dependent SPARC upregulation in head and neck cancer.

## Methods

### Cell culture

Human head and neck cancer cell lines BICR10 and SCC9 were obtained from European Collection of Authenticated Cell Cultures (ECACC). HEK293T cell line was obtained from American Type Cell Culture (ATCC). Cells were cultured in media recommended by ECACC and ATCC, supplemented with 10% FBS (Sigma, F7524), 100 units/mL of penicillin and streptomycin (Invitrogen, 15140122) and 4 mmol/L L-glutamine (Sigma, G7513). All cell lines were cultured in a humidified atmosphere at 37°C and 5% CO_2_. Transfection of overexpression- and promoter-deletion-constructs was performed using polyethylenimine (PEI) (Polysciences, 23966). For crotonate supplementation, cells were treated with 2.5 mM sodium crotonate for 36 h before harvesting.

### Head and neck cancer sample collection

The study was approved by the Institute Ethics Committee of the Indian Institute of Science Education and Research Bhopal, India. Head and neck cancer samples were collected after obtaining informed consent from patients undergoing surgery at Bansal Hospital, Bhopal, India. Details of patients are given in **Supplementary Table 6**.

### Bioinformatics analyses

In order to study cancer associated epigenetic factors we analyzed The Cancer Genome Atlas (TCGA) mRNA expression dataset using TCGAbiolinks package ^37^ in R to obtain epigenetics-related genes that are significantly upregulated in head and neck cancer as compared to normal tissues. Survival analysis was then performed using an online tool OncoLnc (http://www.oncolnc.org). Results of this analysis are given in **Supplementary Table 1**. To obtain the expression of individual genes in TCGA, online platforms like Gepia2 ^38^ and UALCAN ^39^. Correlation analysis was done using Gepia2; EMT hallmark signature (EPITHELIAL_MESENCHYMAL_TRANSITION) used was downloaded from MSigDb database ^40^. GSEA was performed as described previously ^41^. Stratification of TCGA gene expression data on the basis of YEATS2 expression was done and the enrichment of various metabolic pathways^18^ in the YEATS2_high vs. YEATS2_low group was performed using GSEA software ^42^. Results of this analysis are provided in **Supplementary Table 3**.

### RNA interference

SCC9 and BICR10 cells (2 × 10^5^) were seeded in a 6-well plate. After 24 h of seeding, cells were infected with lentivirus containing small hairpin RNA (shRNA) (Sigma, Mission Human Genome shRNA Library) against *YEATS2*, *SP1, EP300* and *GCDH* or shControl plasmid with 8 μg/mL polybrene (Sigma, H9268) containing media. Selection was then done using puromycin at 1 μg/mL (Sigma, P9620) for 3 days before harvesting. Sequences of shRNA are provided in **Supplementary Table 8**.

### Molecular cloning

The *YEATS2* gene was cloned in pEGFP-C3, whereas *SPARC, SP1,* and *GCDH* were cloned in pCMV-3Tag-1A overexpression plasmid (amplified using Phusion Polymerase, NEB, M0530S). *YEATS2* was cloned between SalI and SmaI sites, *SPARC* between BamHI and HindIII sites, *SP1* between EcoRI and SalI sites, and *GCDH* between HindIII and SalI sites. The construct was then confirmed using colony PCR and Sanger sequencing. Promoter deletion constructs of YEATS2 were cloned in pGL3-Basic vector between KpnI and HindIII RE sites. cDNA or genomic DNA from BICR10 cells was used as template in all PCR reactions.

### Site directed mutagenesis

The site-directed mutant construct of the YEATS2 promoter-deletion construct was prepared by using a strategy previously described ^43^. Oligonucleotides with mutations in the SP1-binding sites present from -71 bp to -60 bp upstream of the YEATS2 TSS (cc<u>CGC</u>cc<u>CGC</u>cc<u>CGC</u>cct to cc<u>ATA</u>cc<u>ATA</u>cc<u>ATA</u>cct) were used. The wild-type YEATS2 Luc-508 promoter-deletion construct was used as a template. The primers used are given in **Supplementary Table 7**. The mutation was confirmed by DNA Sanger sequencing.

### Luciferase assay

BICR10 and SCC9 cells (2.5 ×10^5^) were seeded in 6 well plate. After 24 h, cells were co-transfected with either YEATS2-promoter deletion wild-type constructs or SP1-site mutated constructs and pRL-TK Renilla (Promega, E2231) luciferase plasmid. After 24 h of transfection cells were lysed using passive lysis buffer (1% Triton X-100, 25mM Tricine pH 7.8, 15mM potassium phosphate pH 7.8, 15mM magnesium sulphate, 4mM EGTA, 1mM DTT). The Firefly/ Renilla luciferase activity was measured using a luminescence detecting plate reader (CYTATION 5, BioTeK).

### Transwell invasion assay

BICR10 and SCC9 (2 × 10 ^4^) cells resuspended in FBS-free media were added to the upper chamber of transwell setup over a Matrigel layer (Corning, 356230) and incubated for 24 h in a cell culture incubator. Lower chamber of the setup contained media supplemented with FBS. The non-migrated cells in the upper layer of Matrigel were removed, and cells migrated to the lower chamber of transwell were fixed in 4% formaldehyde and then stained with crystal violet (0.05% in 10% methanol in 1 × PBS). Five random fields were counted using an inverted microscope (Olympus CKX41).

### Wound healing assay

BICR10 and SCC9 (1.2 x 10^5^) cells were seeded in a 12-well plate. 12 h after the cells had attached, a scratch was made in the center of each well and wound healing was monitored for 2 days. Three random regions per well were monitored for migration using EVOS FL Auto 2 Imaging System by Thermo Scientific.

### 3D invasion assay

BICR10 and SCC9 (1 x 10^4^) cells were seeded in 96 Well Round (U) Bottom Plate. 48 hours after cell seeding, spheres were transferred to a different flat bottom 96 well plate coated with collagen matrix (Rat Tail Collagen I, Corning 354236) and invasion was checked after 4 days. Images were taken using Olympus CKX41 microscope.

### Immunoblotting

Cells were lysed using urea lysis buffer (8M urea, 2M thiourea, 2% CHAPS, 1% DTT) supplemented with 1× protease inhibitor cocktail (0.5 mM Leupeptin, 250 μM pepstatin, 50 mM EDTA, 1 mM PMSF) at 4°C for 30 min and spun at maximum speed in a 4°C centrifuge for 1 hr. Equal amount of samples were loaded on denaturing polyacrylamide gel and transfer was done to activated PVDF membrane. After transfer, the blots were incubated with recommended dilutions of primary antibodies overnight at 4°C, followed by 40 min incubation with secondary antibody. The blots were scanned using Odyssey Membrane Scanning System. Antibodies used: YEATS2 (Proteintech, 24717-1-AP), SP1 (CST, 9389S), SPARC (CST, 5420S), GCDH (Sigma, HPA043252), N-Cadherin (Abcam, ab19348), Vimentin (Abcam, ab137321), Twist1 (CST, 46702S), GAPDH (CST, 5174S), GFP (Affinity Biosciences, T0006), Flag-tag (Novus, NBP1-06712SS), H3K27cr (PTM Bio, PTM-545RM), Histone H3 (Active Motif, 61475), p300 (CST, 54062S).

### Co-immunoprecipitation

Cells were lysed in Co-IP lysis buffer (50 mM Tris HCl pH 7.5, 150 mM NaCl, 1 mM EDTA, 10% glycerol, 1% NP-40) at 4°C for 1 h. Lysate was spun at maximum speed at 4°C for 1 h in a table-top centrifuge. Co-IP mix for p300 IP and control IP was prepared by taking 1 mg of lysate and diluting it with Co-IP lysis buffer to make a total volume of 500 μl. Antibody for IP (p300, CST:54062S) and control IP (Normal Rabbit IgG, CST: 2729S) was added to the respective tubes and incubated overnight at 4°C. After incubation, 15 μl of Protein G Dynabeads (Invitrogen, 10004D) was added to each tube and incubated for 2 h at 4°C, followed by washing using Co-IP lysis buffer. The captured protein complexes were eluted by adding Laemmli buffer to the beads. The tubes containing the beads were then heated at 95°C for 5 min. The eluant was loaded on SDS-PAGE gel and immunoblotting was performed using standard protocol.

### Immunoflorescence

Cells were plated at a seeding density of 2 × 10^5^ on a coverslip in a 6-well plate. Next day, cells were washed thrice with ice-cold PBS followed by 4% formaldehyde fixation and permeabilization with 0.1% Triton X-100 for 15 min. Blocking was done for 1 h at RT using 2% BSA solution. Cells were then incubated with primary antibodies overnight at 4°C (H3K27cr at 1:200; GCDH at 1:50). Cells were washed thrice with PBS and incubated with Alexa-Flour 555 anti-rabbit IgG secondary antibody for 1 h at RT. The cells were then counterstained with Hoechst 33342 (Invitrogen, HI399) and mounted using fluoroshield (Sigma, F6182). Imaging was performed using Olympus FV3000 live cell microscope with a 60× and 100× oil objective lenses, and image analysis was done using Image J software.

### Immunohistochemistry

Formalin-fixed, paraffin-embedded human head and neck cancer tissue sections were obtained from Bansal Hospital, Bhopal, India. Clinical characteristics of patients used in the study are presented in **Supplementary Table 6**. As described previously ^44^, slides were kept at 65°C for 2 h on hot plate, deparaffinized and, rehydrated as described previously. Heat induced antigen retrieval was then performed using 10 mM sodium citrate buffer (pH 6 in microwave for 14 min). 1:10 dilution of 3% hydrogen peroxidase in methanol was used to quench endogenous peroxidase. Blocking was done with 3% BSA. Primary antibodies against ECHS1 (Sigma, HPA022476; 1:50 dilution), H3K27cr (1:200), GCDH (1:50), YEATS2 (1:50), and Twist1 (1:50) were used. HRP/DAB-chromogenic based Super Sensitive Polymer-HRP Detection System kit (BioGenex, QD430-XAKE) was used as per manufactures’ instructions for staining. All slides were counterstained with Harris’ hematoxylin (Merck). The images were captured by Thermo Scientific Invitrogen EVOS FL Auto 2 Imaging System. Images were then processed in Adobe Photoshop Version 7.0.

### Chromatin immunoprecipitation

ChIP assay was performed as described previously ^45^. Briefly, cells were fixed with 1% formaldehyde and quenched by 0.125 M glycine followed by cell lysis. Further, cross-linked chromatin was sonicated to a chromatin fragment length of 100–400 bp approximately and chromatin was immunoprecipitated using YEATS2, H3K27cr, H3K27ac, SP1 and p300 antibody or control (immunoglobulin G) antibody overnight at 4°C followed by addition of Protein G Dynabeads (Invitrogen, 10004D). The obtained complex and input DNA were de-crosslinked in TE buffer with 1% SDS and proteinase K (Invitrogen, 25530049) at 65°C for overnight. Eluted DNA was then purified using PCR purification kit (QIAGEN, 28106). The immunoprecipitated (IP) DNA and 5% input were analyzed by qPCR using locus specific primers (**Supplementary Table 7**) and SYBR Green Master Mix (Promega, A6002). IP DNA values were normalized to input using the following formula: 2^ (Ct_input -Ct_IP). Resultant values were subsequently normalized to IgG control IP DNA values. The relative values are represented as mean ± SEM of triplicates.

### RT-qPCR

Total was extracted using TRIzol (Invitrogen, 15596026) according to the manufacturer’s instructions. cDNA was synthesized from 3 μg of total RNA by PrimeScript 1st strand cDNA Synthesis Kit (TaKaRa, 6110A). Amplification reactions were performed on light cycler 480 II (Roche) using Go tag SYBR Green master mix (Promega, 75665). The gene expression was calculated using the following formula: 2^ (Ct_control -Ct_target) where RPS16 is taken as control. Sequence of each primer used is given in **Supplementary Table 7**.

### RNA-seq

RNA from shControl/ sh*YEATS2* BICR10 cells was extracted using TRIzol. QIAseq Stranded mRNA Lib Kit (Qiagen, 180441) kit was used to prepare libraries. Paired end sequencing (2×150 bp) of these libraries was performed on Novaseq 6000 platform. Fastq files obtained after sequencing were aligned to GRCh38 human genome using STAR aligner ^46^. Read counting for each sample was done using HTseq2 ^47^. Differential expression analysis was then performed using DESeq2 ^48^. Genes with significant change (FDR corrected p-value < 0.05) in expression are listed in **Supplementary Table 2**.

### ChIP-seq

Eluted DNA obtained after performing ChIP assay in BICR10 cells using YEATS2 antibody. TruSeq ChIP Library Preparation Kit (IP-202-1012) was used to prepare libraries. Paired end sequencing (2^100 bp) of these libraries was performed on Novaseq 6000 platform. Fastq files obtained after sequencing were aligned to GRCh38 human genome using STAR aligner. Peak calling was performed using MACS2 with ‘--keep-dup=all’ parameter ^49^. Genomic distribution of YEATS2 peaks was obtained using ChIPSeeker. For visualizing YEATS2 ChIP-seq signal over *SPARC* promoter, both replicate BAM files were merged using ‘samtools merge’ ^50^ and then bigWig file was generated using ‘bamCoverage’ function of deepTools ^51^. Genes having YEATS2 peak in their promoter region were obtained using Biomart by Ensembl (listed in **Supplementary Table 4**). For H3K27cr ChIP-seq, ChIP assay was performed using chromatin from shControl and sh*YEATS2* BICR10 cells. Libraries were made using NEBNext Ultra II DNA library preparation for Illumina Kit (NEBNext, E7645S) and paired end sequencing (2^150 bp) was performed on Novaseq 6000 platform. Fastq files were aligned to hg38 genome using Bowtie2 ^52^, followed by peak calling with MACS2 and differential binding analysis using DiffBind^53^. Differential sites with p-value < 0.05 were considered significant.

### Statistical analysis

Data are presented as mean ± SEM unless otherwise stated. At least three independent biological replicates have been performed for each experiment. All statistical analyses were conducted using GraphPad Prism 8 software. Statistical tests used and specific *p* values are indicated in the figure legends.

## Acknowledgements

D.P. is a recipient of funding from University Grants Commission, India. P.K. is a recipient of post-doctoral fellowship from Indian Institute of Science Education and Research Bhopal, India, and Department of Biotechnology (DBT, India), Research Associate fellowship award. This work is funded by a grant from the Science and Engineering Research Board (SERB) (STR/2020/000093) awarded to S.S.

## Data availability

YEATS2 and H3K27cr ChIP-Seq data is deposited at GEO database (GSE275975). RNA sequencing data from shControl and sh*YEATS2* cells has been deposited at GEO database (GSE275977). This study does not report any original code.

## Author contributions

S.S. and D.P. conceived and designed the study. D.P., P.K., R.J. and A.S. performed the experiments and analysed the results. D.P. wrote the manuscript with inputs from all other authors. S.S. supervised the experiments and manuscript preparation. S.S. acquired funding for the study.

## References

1. Smith, A., Teknos, T. N. & Pan, Q. Epithelial to mesenchymal transition in head and neck squamous cell carcinoma. Oral Oncol 49, 287–292 (2013).

2. Ribatti, D., Tamma, R. & Annese, T. Epithelial-Mesenchymal Transition in Cancer: A Historical Overview. Transl Oncol 13, 100773 (2020).

3. Sharma, S., Kelly, T. K. & Jones, P. A. Epigenetics in cancer. Carcinogenesis 31, 27–36 (2010).

4. Zhao, S., Allis, C. D. & Wang, G. G. The language of chromatin modification in human cancers. Nat Rev Cancer 21, 413–430 (2021).

5. Wellen, K. E. et al. ATP-Citrate Lyase Links Cellular Metabolism to Histone Acetylation. Science (1979) 324, 1076–1080 (2009).

6. Wang, Y. et al. KAT2A coupled with the α-KGDH complex acts as a histone H3 succinyltransferase. Nature 552, 273–277 (2017).

7. Pandkar, M. R. et al. PKM2 dictates the poised chromatin state of PFKFB3 promoter to enhance breast cancer progression. NAR Cancer 5, zcad032 (2023).

8. Diehl, K. L. & Muir, T. W. Chromatin as a key consumer in the metabolite economy. Nat Chem Biol 16, 620–629 (2020).

9. Zhao, D. et al. YEATS2 is a selective histone crotonylation reader. Cell Res 26, 629–632 (2016).

10. Liao, M. et al. LINC00922 decoys SIRT3 to facilitate the metastasis of colorectal cancer through up-regulation the H3K27 crotonylation of ETS1 promoter. Mol Cancer 22, 163 (2023).

11. Santos, C. R. et al. VRK1 Signaling Pathway in the Context of the Proliferation Phenotype in Head and Neck Squamous Cell Carcinoma. Molecular Cancer Research 4, 177–185 (2006).

12. Johnson, J. J. et al. Protease-activated Receptor-2 (PAR-2)-mediated Nf-κB Activation Suppresses Inflammation-associated Tumor Suppressor MicroRNAs in Oral Squamous Cell Carcinoma*. Journal of Biological Chemistry 291, 6936–6945 (2016).

13. Xie, G. et al. The role of imprinting genes’ loss of imprints in cancers and their clinical implications. Front Oncol Volume 14-2024, (2024).

14. Zhang, H., Chen, W., Fu, X., Su, X. & Yang, A. CBX3 promotes tumor proliferation by regulating G1/S phase via p21 downregulation and associates with poor prognosis in tongue squamous cell carcinoma. Gene 654, 49–56 (2018).

15. Liu, Y. et al. High expression of ACTL6A leads to poor prognosis of oral squamous cell carcinoma patients through promoting malignant progression. Head Neck 46, 1450–1467 (2024).

16. Li, W. et al. BOP1 Used as a Novel Prognostic Marker and Correlated with Tumor Microenvironment in Pan-Cancer. J Oncol 2021, 3603030 (2021).

17. Taniue, K. et al. RNA Exosome Component EXOSC4 Amplified in Multiple Cancer Types Is Required for the Cancer Cell Survival. Int J Mol Sci 23, (2022).

18. Su, A. et al. The Folate Cycle Enzyme MTHFR Is a Critical Regulator of Cell Response to MYC-Targeting Therapies. Cancer Discov 10, 1894–1911 (2020).

19. Leandro, J. & Houten, S. M. The lysine degradation pathway: Subcellular compartmentalization and enzyme deficiencies. Mol Genet Metab 131, 14–22 (2020).

20. Rivera, L. B., Bradshaw, A. D. & Brekken, R. A. The regulatory function of SPARC in vascular biology. Cellular and Molecular Life Sciences 68, 3165–3173 (2011).

21. Tremble, P. M., Lane, T. F., Sage, E. H. & Werb, Z. SPARC, a secreted protein associated with morphogenesis and tissue remodeling, induces expression of metalloproteinases in fibroblasts through a novel extracellular matrix-dependent pathway. Journal of Cell Biology 121, 1433–1444 (1993).

22. Xie, J. et al. The mechanisms, regulations, and functions of histone lysine crotonylation. Cell Death Discov 10, 66 (2024).

23. Sabari, B. R. et al. Intracellular Crotonyl-CoA Stimulates Transcription through p300-Catalyzed Histone Crotonylation. Mol Cell 58, 203–215 (2015).

24. Kinnaird, A., Zhao, S., Wellen, K. E. & Michelakis, E. D. Metabolic control of epigenetics in cancer. Nat Rev Cancer 16, 694–707 (2016).

25. Leshem, O. et al. TMPRSS2/ERG Promotes Epithelial to Mesenchymal Transition through the ZEB1/ZEB2 Axis in a Prostate Cancer Model. PLoS One 6, e21650-(2011).

26. Yuxiong, W. et al. Regulatory mechanisms of the cAMP-responsive element binding protein 3 (CREB3) family in cancers. Biomedicine & Pharmacotherapy 166, 115335 (2023).

27. Zhao, D., Li, Y., Xiong, X., Chen, Z. & Li, H. YEATS Domain—A Histone Acylation Reader in Health and Disease. J Mol Biol 429, 1994–2002 (2017).

28. Liu, N. et al. Histone H3 lysine 27 crotonylation mediates gene transcriptional repression in chromatin. Mol Cell 83, 2206–2221.e11 (2023).

29. Tan, M. et al. Identification of 67 Histone Marks and Histone Lysine Crotonylation as a New Type of Histone Modification. Cell 146, 1016–1028 (2011).

30. Jiang, G., Li, C., Lu, M., Lu, K. & Li, H. Protein lysine crotonylation: past, present, perspective. Cell Death Dis 12, 703 (2021).

31. Ma, J. et al. SPARC enhances 5-FU chemosensitivity in gastric cancer by modulating epithelial-mesenchymal transition and apoptosis. Biochem Biophys Res Commun 558, 134–140 (2021).

32. Carriere, P. et al. Role of SPARC in the epithelial-mesenchymal transition induced by PTHrP in human colon cancer cells. Mol Cell Endocrinol 530, 111253 (2021).

33. Sangaletti, S. et al. Mesenchymal Transition of High-Grade Breast Carcinomas Depends on Extracellular Matrix Control of Myeloid Suppressor Cell Activity. Cell Rep 17, 233–248 (2016).

34. Gao, D. et al. P300 regulates histone crotonylation and preimplantation embryo development. Nat Commun 15, 6418 (2024).

35. Li, X., Egervari, G., Wang, Y., Berger, S. L. & Lu, Z. Regulation of chromatin and gene expression by metabolic enzymes and metabolites. Nat Rev Mol Cell Biol 19, 563–578 (2018).

36. Matsuda, S. et al. Nuclear pyruvate kinase M2 complex serves as a transcriptional coactivator of arylhydrocarbon receptor. Nucleic Acids Res 44, 636–647 (2016).

37. Colaprico, A. et al. TCGAbiolinks: an R/Bioconductor package for integrative analysis of TCGA data. Nucleic Acids Res 44, e71–e71 (2016).

38. Tang, Z., Kang, B., Li, C., Chen, T. & Zhang, Z. GEPIA2: an enhanced web server for large-scale expression profiling and interactive analysis. Nucleic Acids Res 47, W556–W560 (2019).

39. Chandrashekar, D. S. et al. UALCAN: An update to the integrated cancer data analysis platform. Neoplasia 25, 18–27 (2022).

40. Liberzon, A. et al. Molecular signatures database (MSigDB) 3.0. Bioinformatics 27, 1739–1740 (2011).

41. Pant, D., Narayanan, S. P., Vijay, N. & Shukla, S. Hypoxia-induced changes in intragenic DNA methylation correlate with alternative splicing in breast cancer. J Biosci 45, 3 (2020).

42. Subramanian, A. et al. Gene set enrichment analysis: A knowledge-based approach for interpreting genome-wide expression profiles. Proceedings of the National Academy of Sciences 102, 15545–15550 (2005).

43. Zhang, K. et al. A high-efficiency method for site-directed mutagenesis of large plasmids based on large DNA fragment amplification and recombinational ligation. Sci Rep 11, 10454 (2021).

44. Kakani, P. et al. Hypoxia-induced CTCF promotes EMT in breast cancer. Cell Rep 43, 114367 (2024).

45. Singh, S. et al. Intragenic DNA methylation and BORIS-mediated cancer-specific splicing contribute to the Warburg effect. Proceedings of the National Academy of Sciences (2017) doi:10.1073/pnas.1708447114.

46. Dobin, A. et al. STAR: ultrafast universal RNA-seq aligner. Bioinformatics 29, 15–21 (2013).

47. Anders, S., Pyl, P. T. & Huber, W. HTSeq—a Python framework to work with high-throughput sequencing data. Bioinformatics 31, 166–169 (2015).

48. Love, M. I., Huber, W. & Anders, S. Moderated estimation of fold change and dispersion for RNA-seq data with DESeq2. Genome Biol 15, 550 (2014).

49. Zhang, Y. et al. Model-based Analysis of ChIP-Seq (MACS). Genome Biol 9, R137 (2008).

50. Li, H. et al. The Sequence Alignment/Map format and SAMtools. Bioinformatics 25, 2078– 2079 (2009).

51. Ramírez, F., Dündar, F., Diehl, S., Grüning, B. A. & Manke, T. deepTools: a flexible platform for exploring deep-sequencing data. Nucleic Acids Res 42, W187–W191 (2014).

52. Langmead, B. & Salzberg, S. L. Fast gapped-read alignment with Bowtie 2. Nat Methods 9, 357–359 (2012).

53. Stark, R. & Brown, G. D. DiffBind : Differential binding analysis of ChIP-Seq peak data. in (2012).

