## Supplementary Figures for "Interplay of YEATS2 and GCDH regulates histone crotonylation and drives EMT in head and neck cancer"

### Figure Supplements

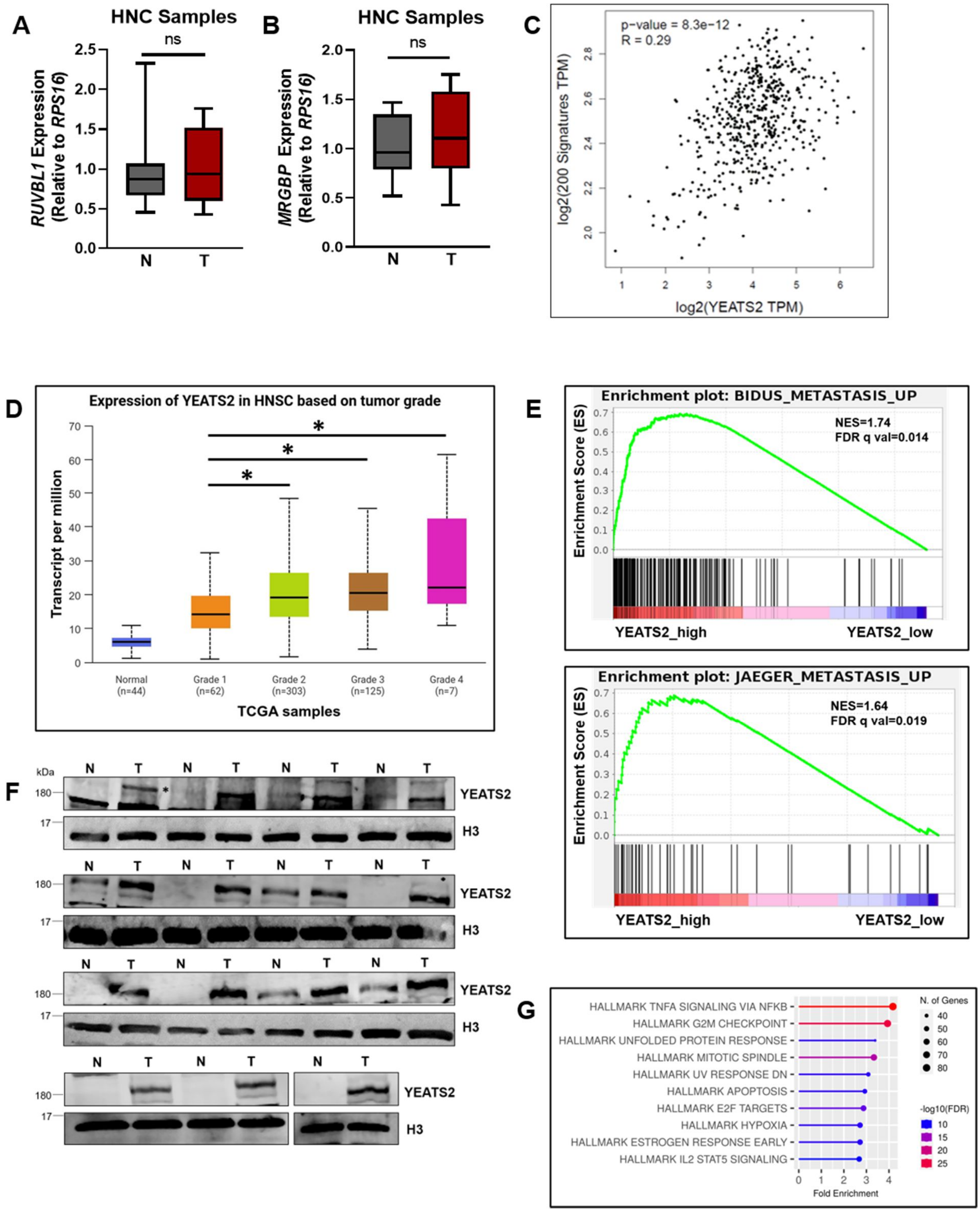


**Figure 1—figure supplement 1. YEATS2 is upregulated in head and neck cancer. (A-B)** RT-qPCR result showing mRNA expression of (A) *RUVBL1* and (B) *MRGBP* in HNC samples. **(C)** Scatter plot showing positive correlation of *YEATS2* expression with the expression of genes included in hallmark

EPITHELIAL_MESENCHYMAL gene signature, in HNC TCGA data. **(D)** Expression of YEATS2 in different grades of TCGA HNC samples, showing significant increase in expression from grade 1 to subsequent grades. **(E)** GSEA plot showing significant enrichment of metastasis-associated gene sets in TCGA samples stratified as YEATS2_high as compared to YEATS2_low samples. **(F)** Immunoblot showing protein levels of YEATS2 in nuclear lysates extracted from HNC samples (YEATS2 band indicated by *). **(G)** Results of overrepresentation analysis of genes significantly upregulated in shControl vs. sh*YEATS2* RNA-seq data. Error bars, min to max; two-tailed t test, ns- non-significant, $*p < 0.05$, N-Normal, T-Tumor.

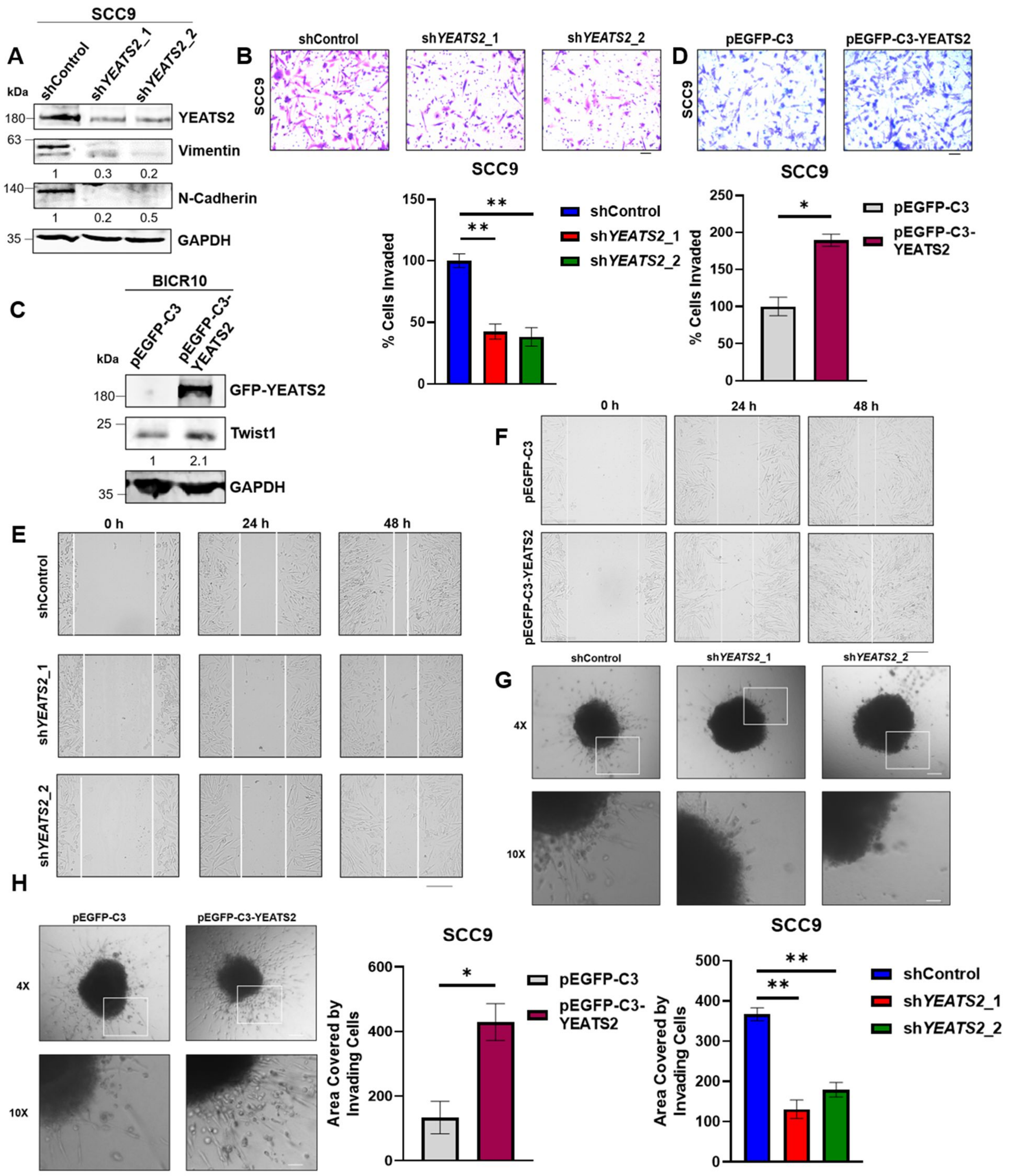


**Figure 2—figure supplement 1. YEATS2 drives EMT in head and neck cancer cells.** **(A)** Immunoblot showing the expression levels of various EMT factors upon YEATS2 knockdown in SCC9. **(B and D)** Results of invasion assay after knockdown (B) or overexpression (D) of YEATS2 in SCC9 (Scale bar, 200 μm). **(C)** Immunoblot showing the expression level of Twist1 upon YEATS2 overexpression in SCC9. **(E-**

**F)** Wound healing assay results performed after knockdown (E) or overexpression (F) of YEATS2 in SCC9 (Scale bar, 275 µm). **(G-H)** Results of 3D invasion assay showing change in invasive potential of SCC9 cells in collagen matrix after silencing (G, quantification below) or overexpression (H, quantification on right) of YEATS2. (Scale bar: 4X, 200 µm; 10X, 50 µm) Error bars, mean ± SEM; two-tailed t test, $*p < 0.05$, $**p < 0.01$, n = 3 biological replicates.

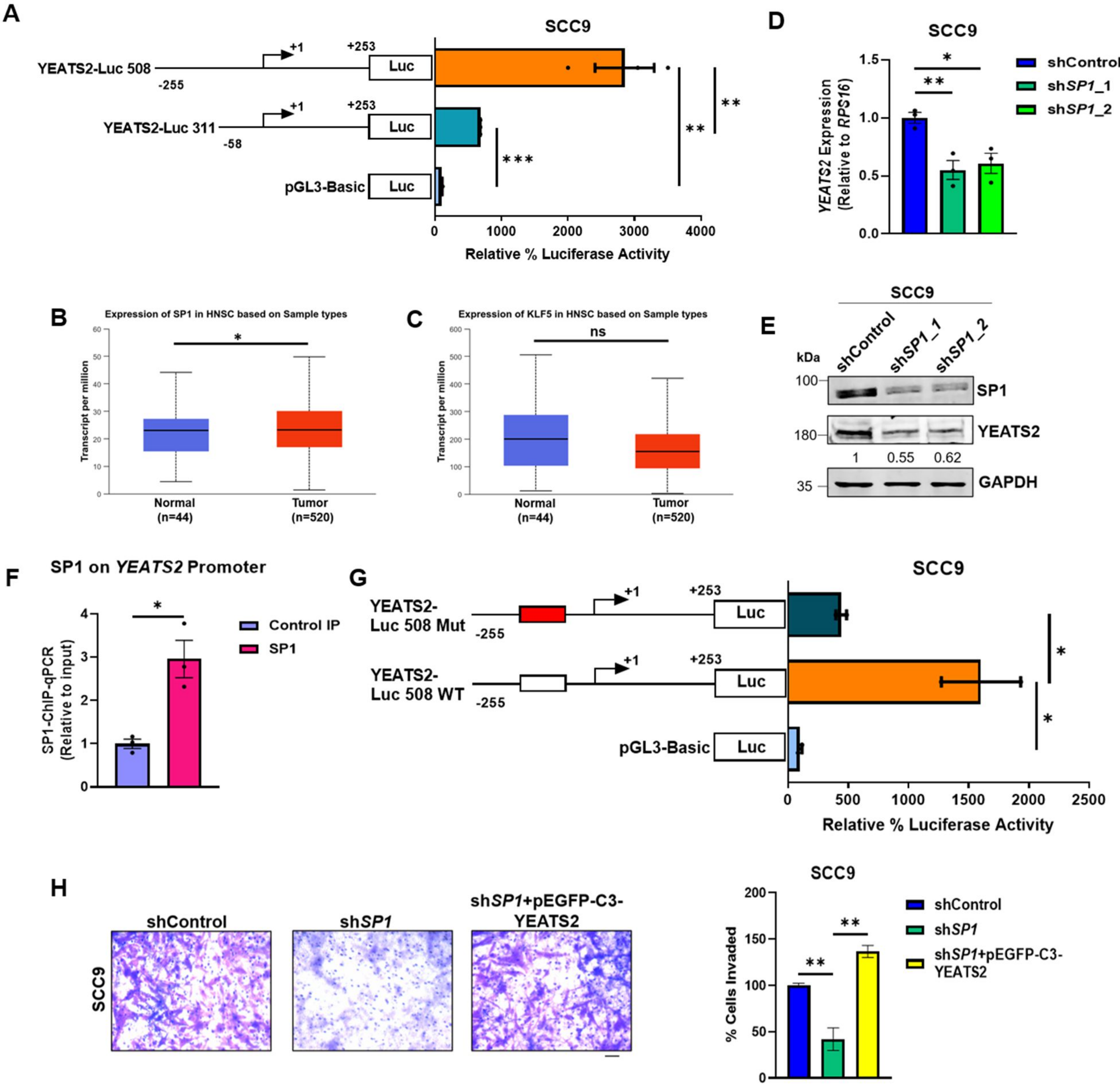


**Figure 3—figure supplement 1. Regulation of EMT by YEATS2 is SP1-dependent in HNC. (A)** Luciferase assay results showing the difference in relative luciferase activity of the two YEATS2 promoter deletion constructs in SCC9. **(B-C)** Expression levels of *SP1* (B) and *KLF5* (C) in TCGA HNC gene expression dataset. **(D)** Plot showing decrease in mRNA expression of *YEATS2* on SP1-knockdown in

SCC9 cells. **(E)** Immunoblot showing the reduced expression level of YEATS2 upon SP1-knockdown in SCC9. **(F)** Plot depicting SP1 binding on *YEATS2* promoter in SP1-ChIP-qPCR assay in SCC9 cells. **(G)** Relative luciferase activity of wild-type (WT) vs. mutant (Mut) YEATS2 Luc-508 in SCC9. **(H)** Invasion assay images (with quantification on right) showing decrease and rescue of the percentage of invaded cells in sh*SP1* SCC9 cells, and sh*SP1* cells with YEATS2 overexpression, respectively. Scale bar, 200 μm. Error bars, mean ± SEM; two-tailed t test, ns- non-significant, $*p < 0.05$, $**p < 0.01$, $***p < 0.001$, n = 3 biological replicates.

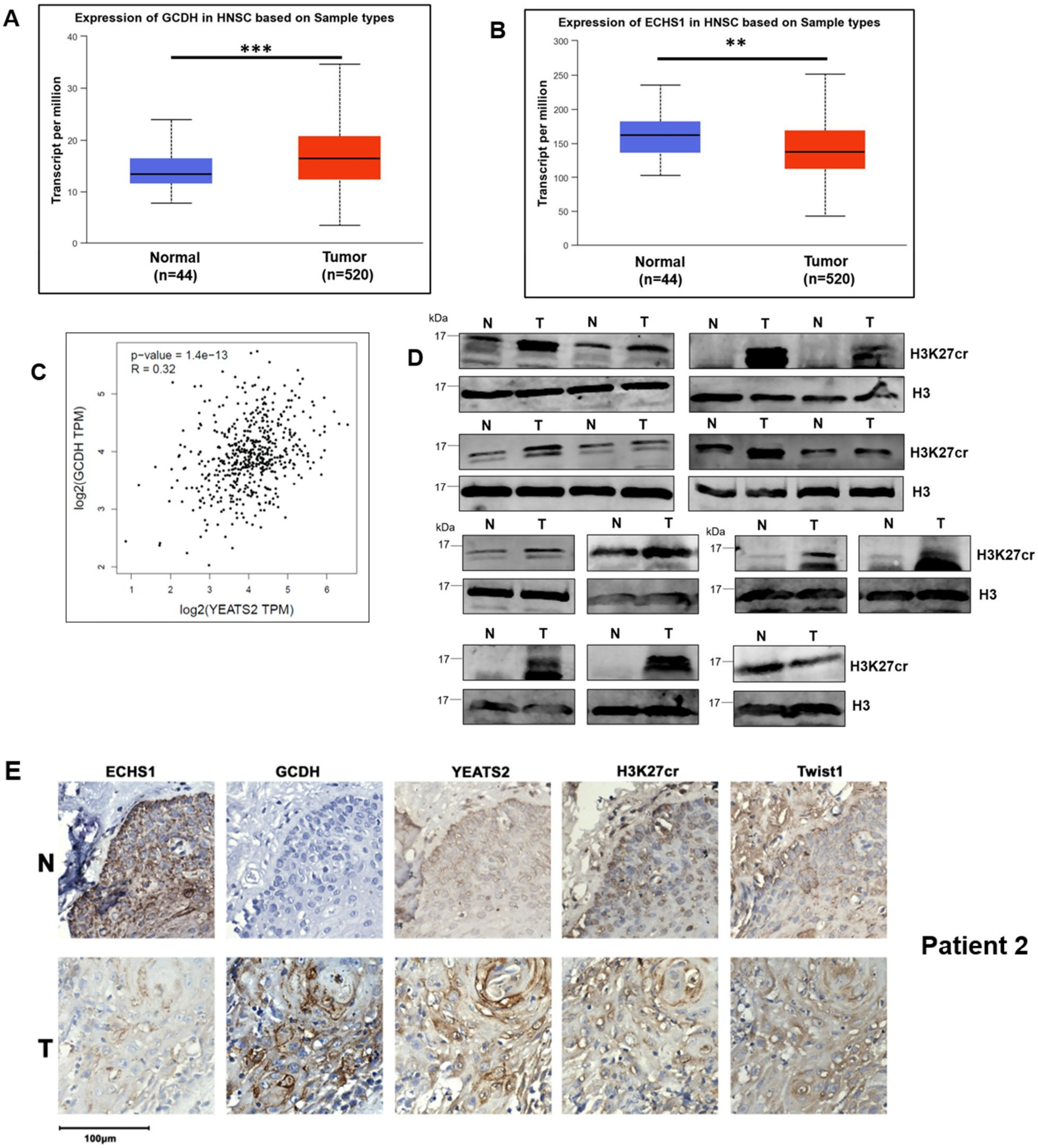

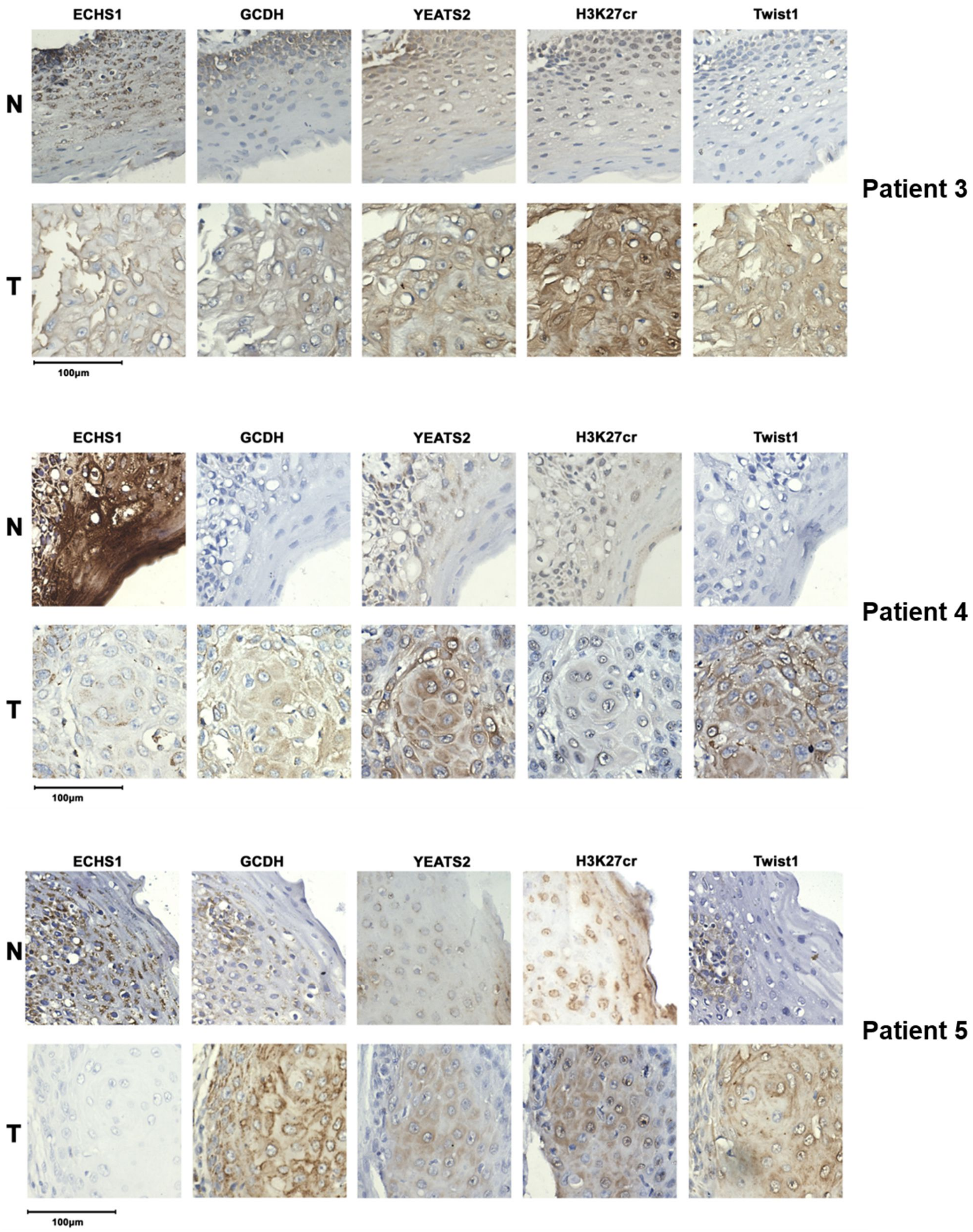
ECHS1
GCDH
YEATS2
H3K27cr
Twist1
N
T
100μm
Patient 3
ECHS1
GCDH
YEATS2
H3K27cr
Twist1
N
T
100μm
Patient 4
ECHS1
GCDH
YEATS2
H3K27cr
Twist1
N
T
100μm
Patient 5

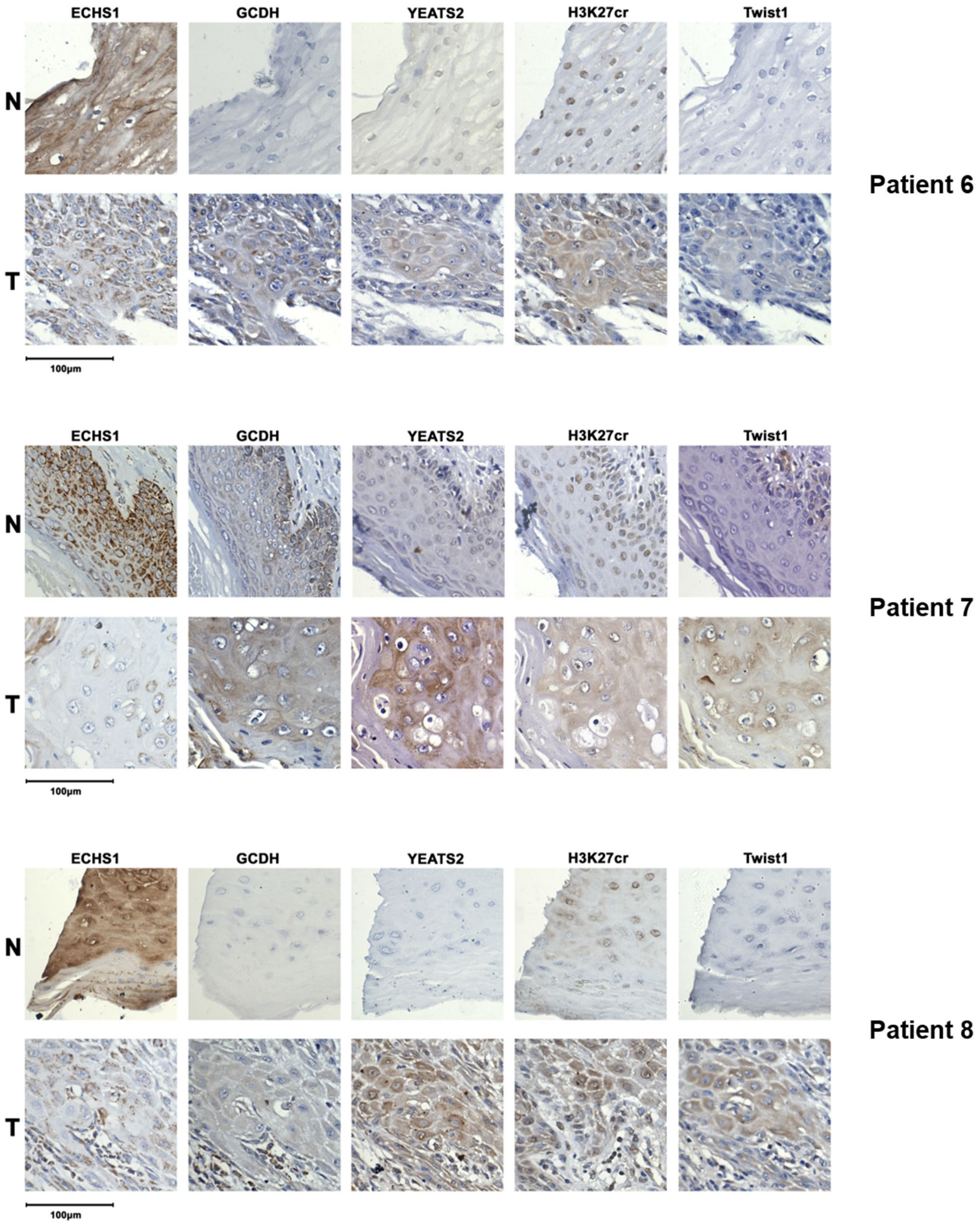
ECHS1
GCDH
YEATS2
H3K27cr
Twist1
N
T
100μm
Patient 6
ECHS1
GCDH
YEATS2
H3K27cr
Twist1
N
T
100μm
Patient 7
ECHS1
GCDH
YEATS2
H3K27cr
Twist1
N
T
100μm
Patient 8

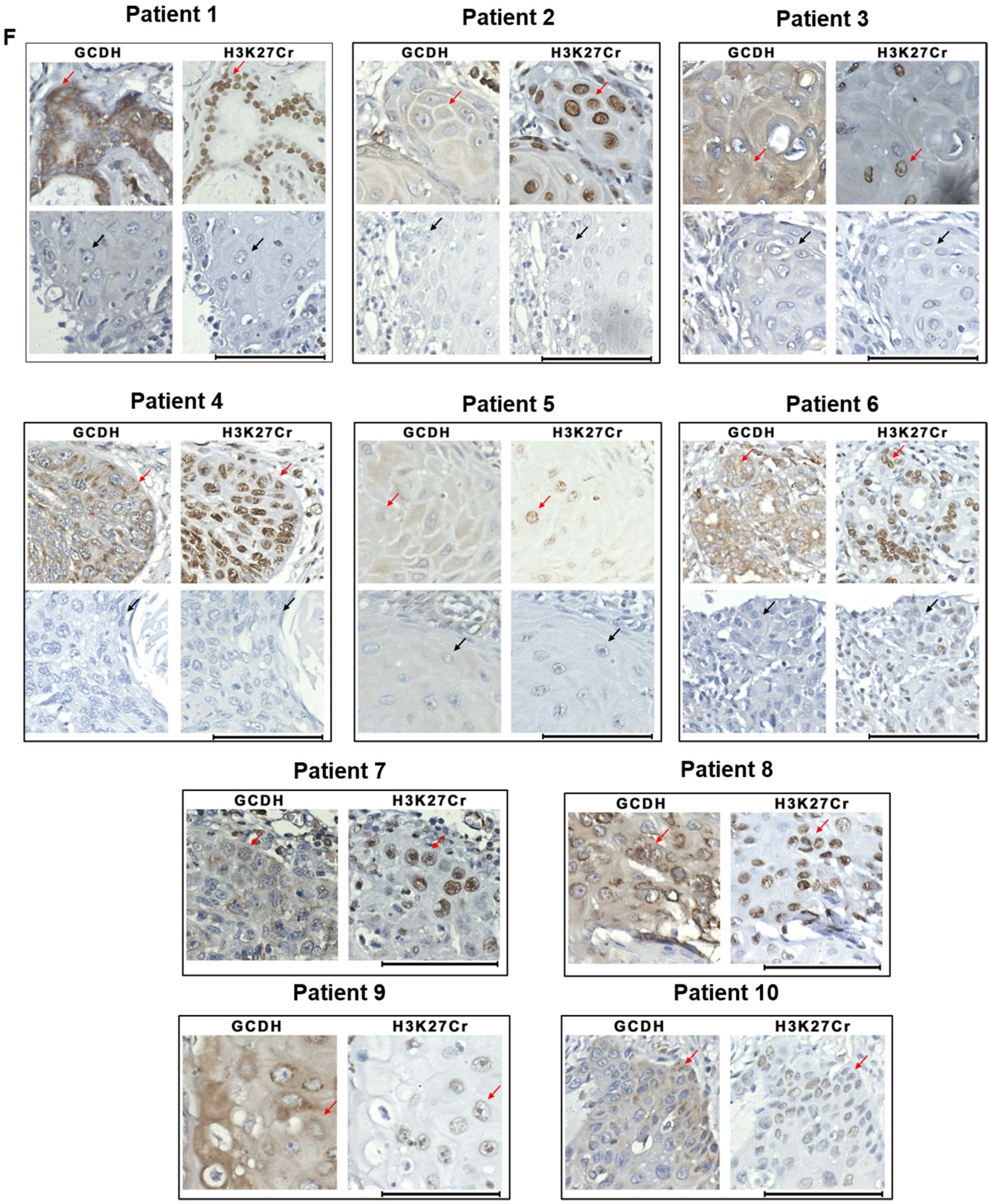
F
Patient 1
GCDH
H3K27Cr
Patient 2
GCDH
H3K27Cr
Patient 3
GCDH
H3K27Cr
Patient 4
GCDH
H3K27Cr
Patient 5
GCDH
H3K27Cr
Patient 6
GCDH
H3K27Cr
Patient 7
GCDH
H3K27Cr
Patient 8
GCDH
H3K27Cr
Patient 9
GCDH
H3K27Cr
Patient 10
GCDH
H3K27Cr

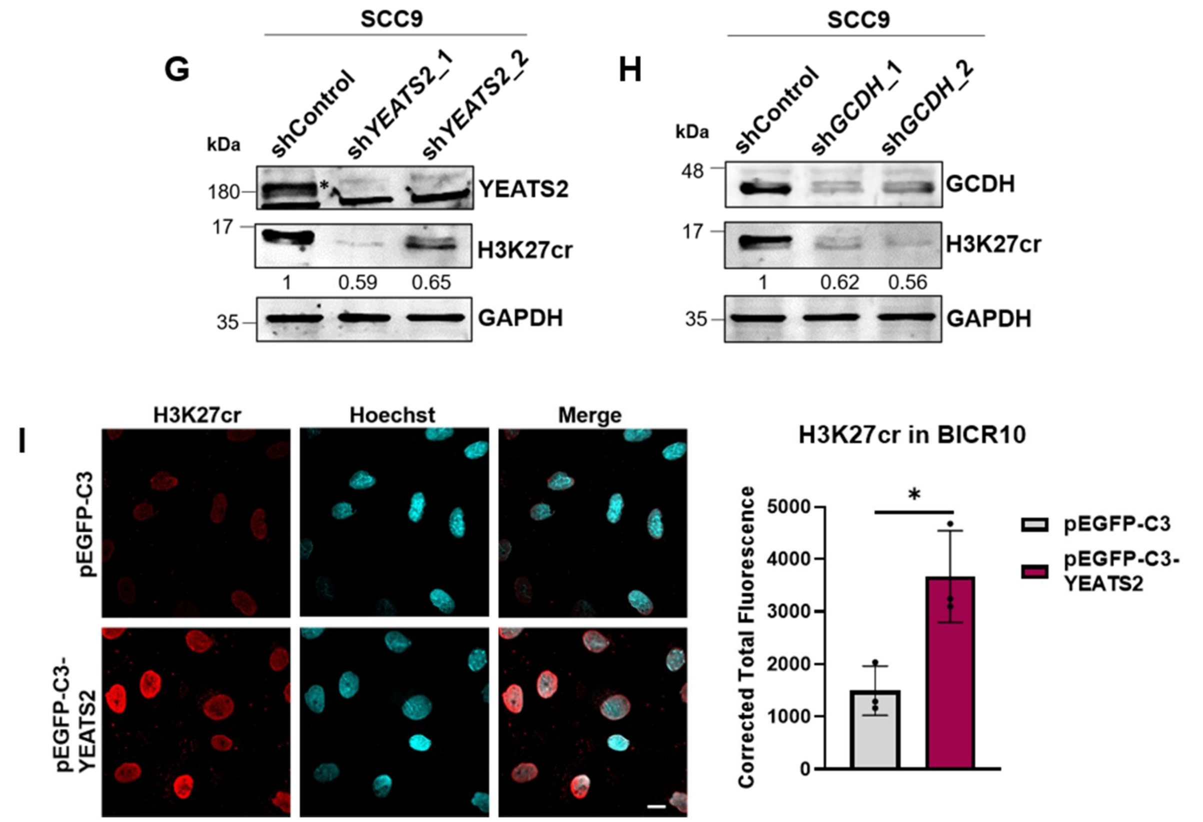


**Figure 4—figure supplement 1. YEATS2 and GCDH regulate histone crotonylation in HNC. (A-B)** Plots showing increase or decrease in expression of *GCDH* (A) and *ECHS1* (B) genes respectively, in HNC TCGA data. **(C)** Scatter plot showing positive correlation between *YEATS2* and *GCDH* expression levels in HNC TCGA data. **(D)** Immunoblot showing enhanced levels of H3K27cr in nuclear lysates extracted from HNC tumor vs. normal samples. **(E)** Representative IHC images showing the levels of ECHS1, GCDH, YEATS2, H3K27cr and Twist1 in HNC normal vs. tumor tissue samples (Scale bar, 100 µm). **(F)** Representative IHC images showing the colocalization of nuclear GCDH with H3K27cr (Scale bar, 100 µm). Cells with nuclear GCDH-H3K27cr localization indicated by red arrows, whereas non-nuclear staining indicated by black arrows. **(G-H)** Immunoblot depicting the decrease in H3K27cr levels on (G) YEATS2- and (H) GCDH-knockdown in SCC9 cells (YEATS2 band indicated by *). **(I)** Immunoflorescence images (quantification on right) depicting the increase in H3K27cr levels on YEATS2 overexpression (Scale bar, 10 µm).

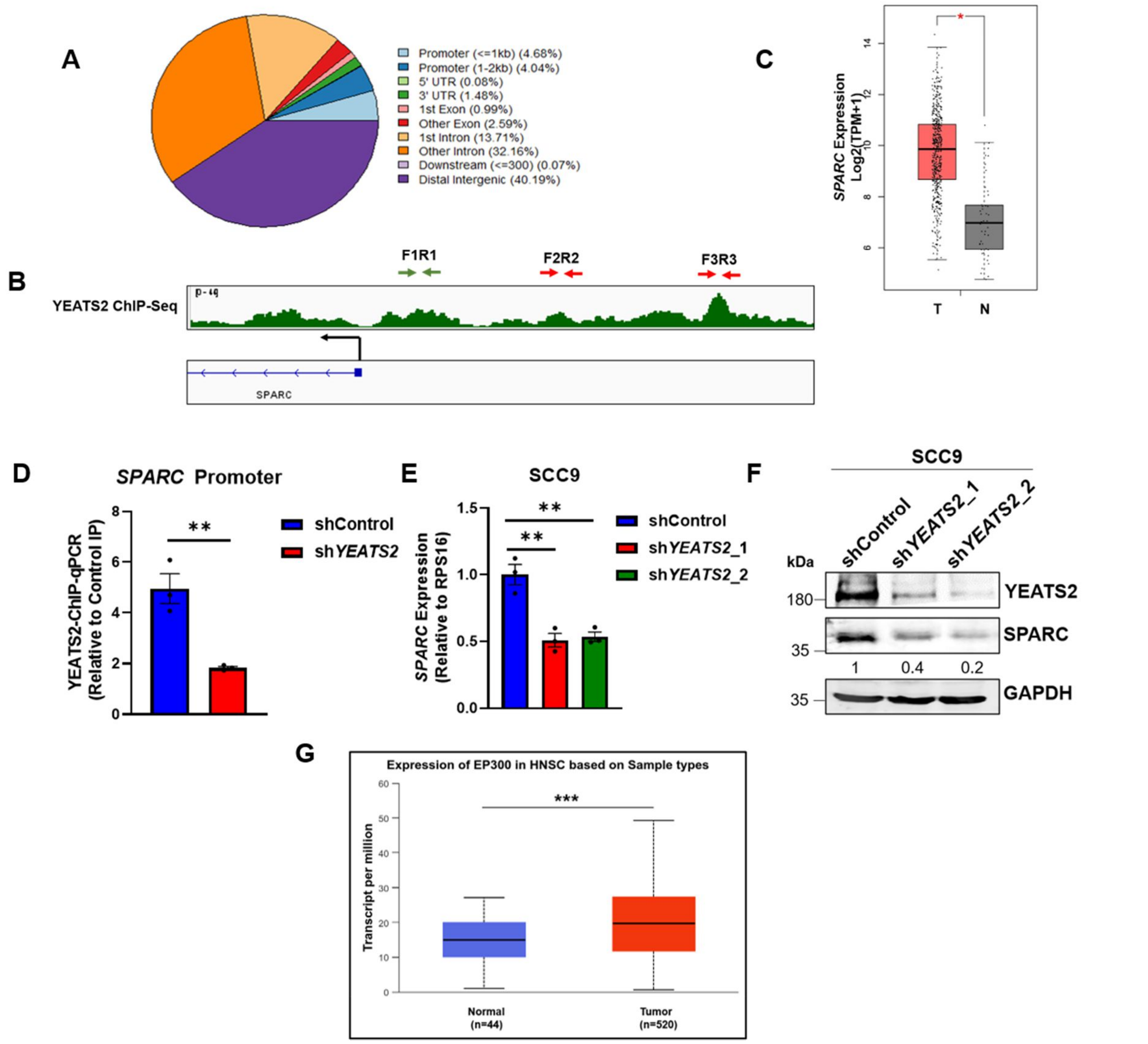


**Figure 5—figure supplement 1. YEATS2 regulates expression of EMT-related SPARC in HNC.** **(A)** Pie chart showing the genomic distribution of YEATS2 peaks in BICR10 cells. **(B)** IGV plot showing the enrichment of YEATS2 on the promoter of *SPARC* in YEATS2 ChIP-seq data. Arrows indicate the regions probed for YEATS2 binding on *SPARC* in YEATS2-ChIP-qPCR assay; region corresponding to F1R1 showed significant change in YEATS2 occupancy in sh*YEATS2* BICR10 cells. **(C)** Increased expression of *SPARC* gene in TCGA HNC tumor vs. normal data. **(D)** YEATS2-ChIP-qPCR results showing decreased binding of YEATS2 on *SPARC* promoter in sh*YEATS2* SCC9 cells. **(E)** RT-qPCR results showing decreased expression of *SPARC* on YEATS2-knockdown in SCC9. **(F)** Immunoblot showing decreased expression of SPARC on YEATS2-knockdown in SCC9. **(G)** *EP300* expression levels in N vs. T HNC samples from TCGA. Error bars, mean ± SEM; two-tailed t test, $**p < 0.01$, $***p < 0.001$, n = 3 biological replicates.

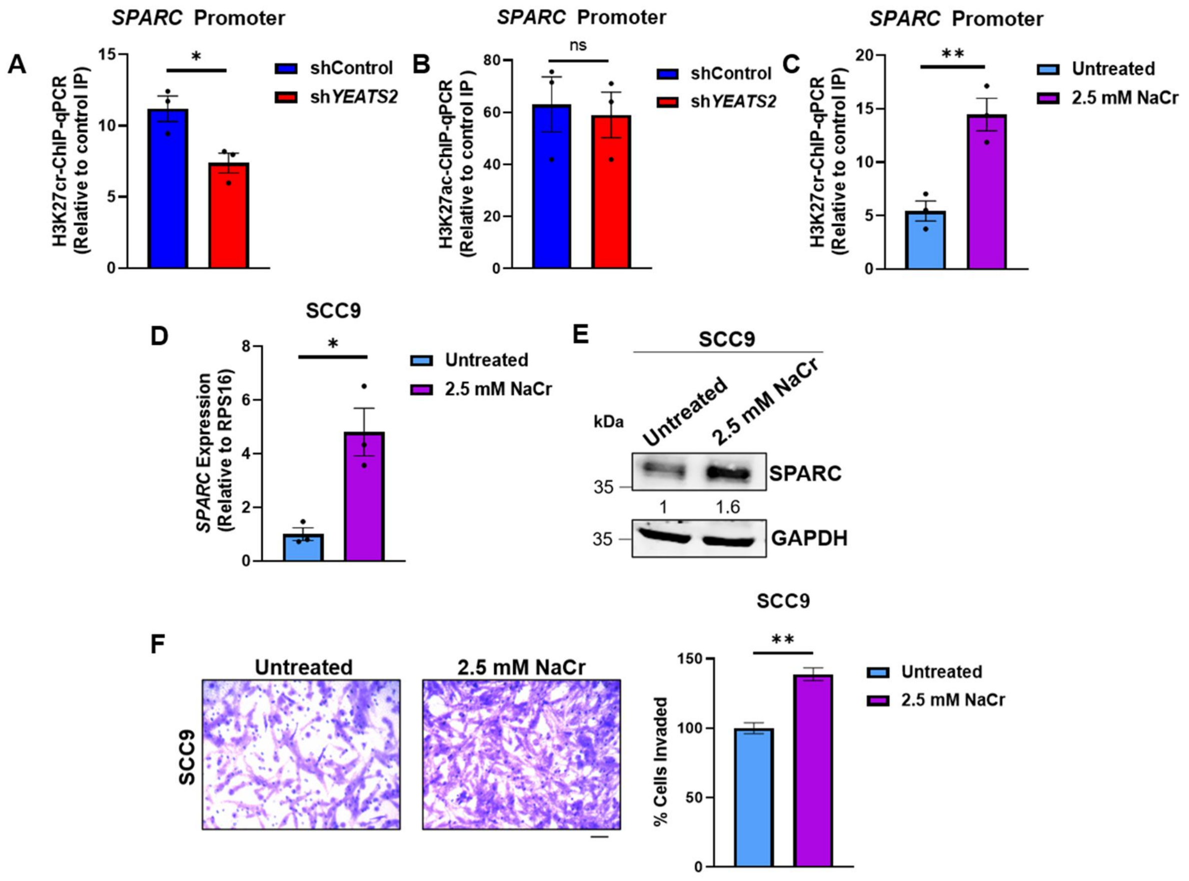


**Figure 6—figure supplement 1. Supplementation of sodium crotonate increases SPARC levels in YEATS2-dependent manner in HNC cells. (A)** H3K27cr-ChIP-qPCR results showing decrease in H3K27Cr enrichment on *SPARC* promoter on YEATS2 knockdown in SCC9 cells. **(B)** H3K27ac-ChIP-qPCR results showing non-significant change in H3K27ac enrichment on *SPARC* promoter on YEATS2 knockdown in SCC9. **(C)** H3K27cr-ChIP-qPCR results showing increase in H3K27cr enrichment on *SPARC* promoter on treating SCC9 cells with 2.5 mM sodium crotonate (NaCr). **(D-E)** RT-qPCR (D) and immunoblot (E) showing enhanced SPARC expression in untreated SCC9 cells vs. SCC9 cells treated with 2.5 mM NaCr. **(F)** Invasion assay images (with quantification on right) showing increased invasion on treating SCC9 cells with 2.5 mM NaCr (Scale bar, 200 μm). Error bars, mean ± SEM; two-tailed t test, ns- non-significant, $*p < 0.05$, $**p < 0.01$.

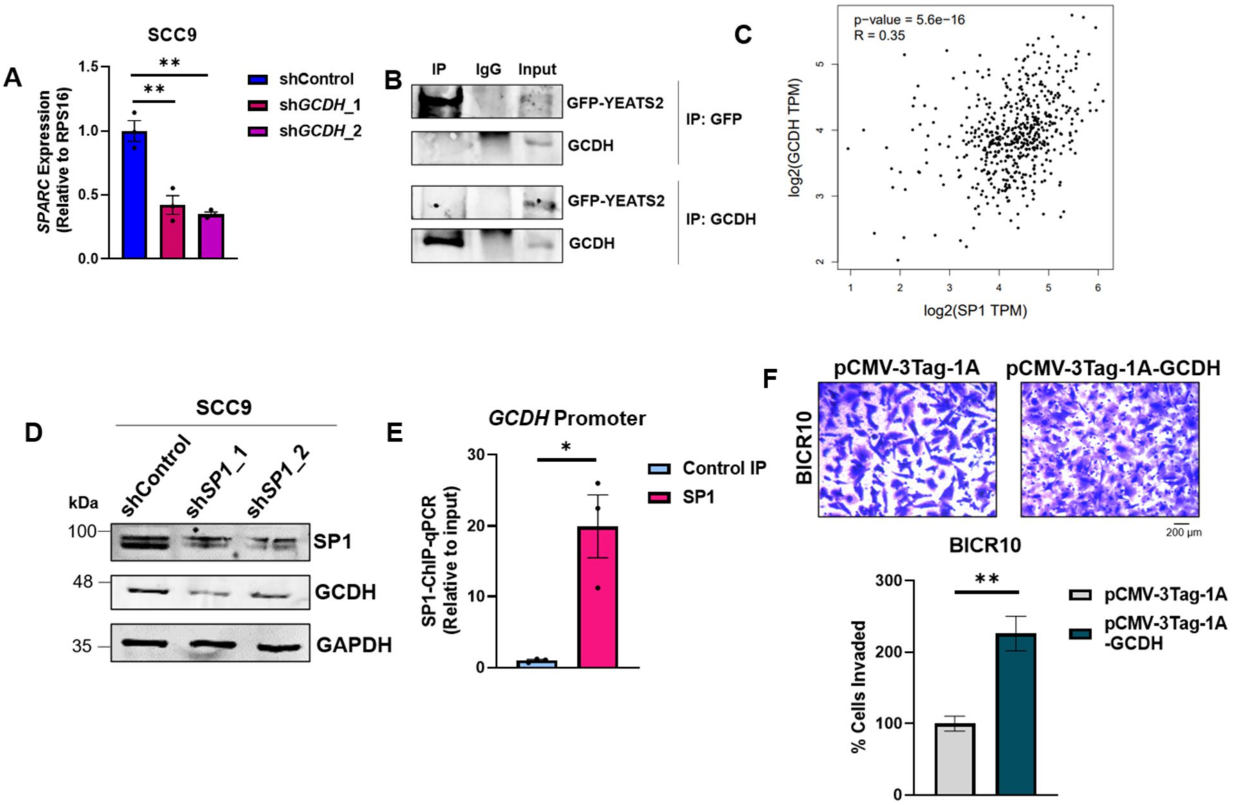


**Figure 7—figure supplement 1. GCDH expression is SP1-dependent and regulates H3K27cr-mediated SPARC expression with YEATS2 synergistically.** **(A)** RT-qPCR results showing decrease in *SPARC* expression on GCDH-knockdown in SCC9 cells. **(B)** Co-IP Immunoblot showing lack of interaction between YEATS2 and GCDH in HEK293T cells. **(C)** Scatter plot showing positive correlation between *GCDH* and *SP1* expression levels in HNC TCGA data. **(D)** Immunoblot showing the reduced expression of GCDH on SP1-knockdown in SCC9 cells. **(E)** Plot showing SP1 binding on *GCDH* promoter in SP1-ChIP assay in SCC9. **(F)** Invasion assay images showing increase in invasion of BICR10 cells on GCDH overexpression (Scale bar, 200 μm). Error bars, mean ± SEM; two-tailed t test, $*p < 0.05$, $**p < 0.01$, n = 3 biological replicates.
